# A conserved in-frame stop codon acts as a multipotent defense mechanism in alphaviruses

**DOI:** 10.1101/2025.09.27.679005

**Authors:** Tamanash Bhattacharya, Tiia S. Freeman, Eva M. Alleman, Fang Wang, Lyuba Chechik, Michael Emerman, Kevin M. Myles, Harmit S. Malik

## Abstract

Most alphaviruses maintain an in-frame opal stop codon that interrupts their non-structural polyprotein (nsP) ORF between nsP3 and nsP4 in both vertebrate and insect hosts. We show that the nsP3 opal stop codon confers a replicative advantage to Sindbis virus (SINV) in RNAi-competent mosquito cells and in *Aedes aegypti* mosquitoes, but not in cells or mosquitoes lacking RNAi. Mutation of the opal stop codon delays processing of the viral nsP polyprotein, disrupts viral replication spherule integrity, and renders viral RNA susceptible to Dicer 2 cleavage, resulting in higher antiviral siRNA responses against SINV. Similarly, these defects caused by opal codon mutations lead to increased viral RNA detection and enhanced immune signaling in vertebrate cells. Thus, a single stop codon in alphaviruses mediates a multipotent viral strategy to evade innate immune defenses across diverse hosts.

**Teaser:** A conserved ORF-interrupting stop codon helps alphaviruses avoid triggering innate antiviral immunity across diverse hosts.

## Introduction

Most alphaviruses encode an in-frame premature opal (UGA) stop codon within the non-structural polyprotein (nsP) open reading frame at the end of the *non-structural protein 3 (nsP3)* gene, just before the *non-structural protein 4 (nsP4)* gene (Figure 1A) (*1*). Programmed ribosomal readthrough (PRT) at this opal stop codon enables alphaviruses to produce two polyproteins from their *nsP* genes: standard translation yields the nsP1-nsP2-nsP3 polyprotein (P123), whereas PRT yields the longer nsP1-nsP2-nsP3-nsP4 polyprotein (P1234) (*2*). Sufficient PRT is essential for viral replication, as nsP4 functions as the viral RNA polymerase required to synthesize new viral RNA (*3*).

**Figure 1.**
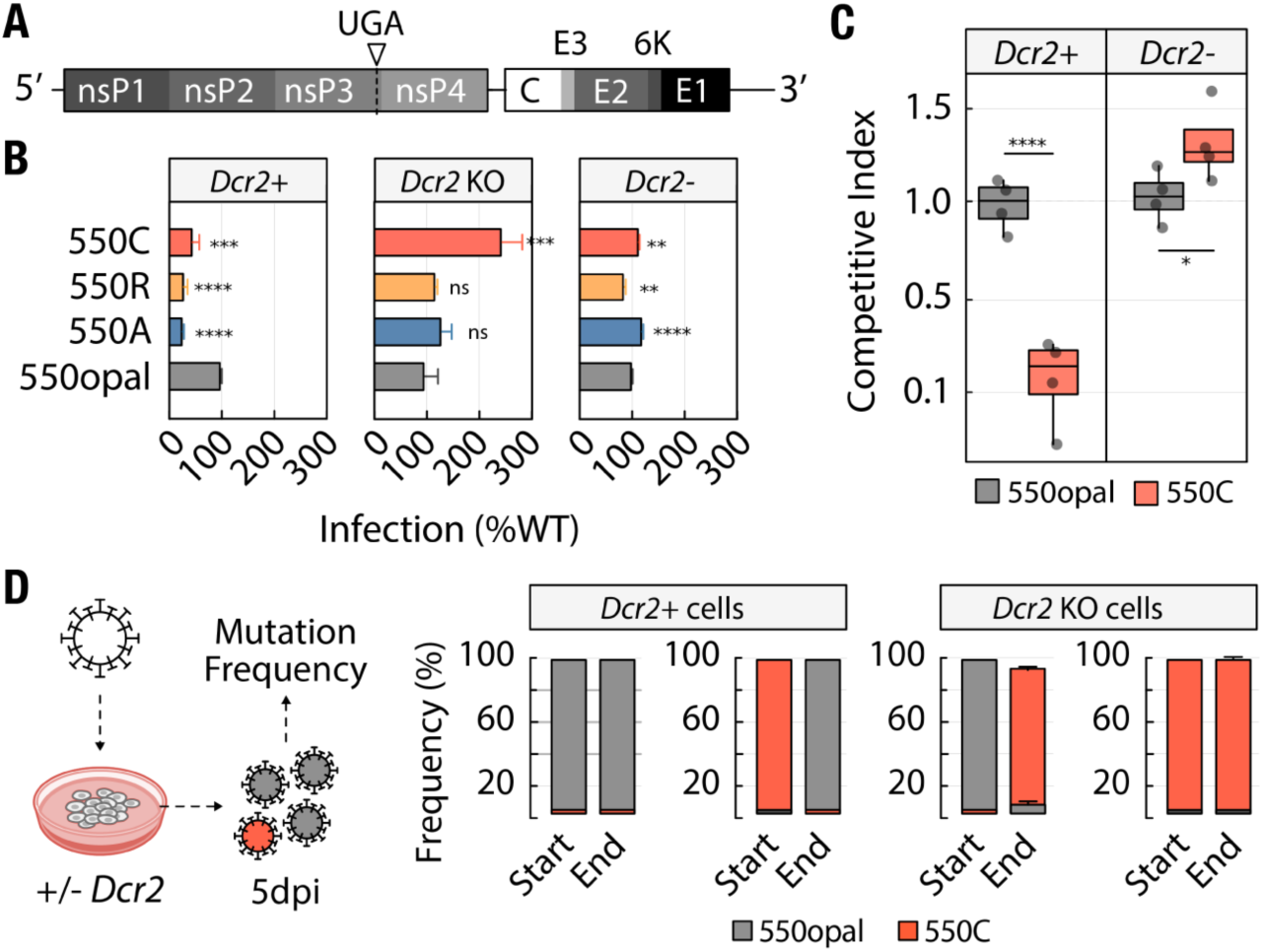
The mosquito RNAi pathway imposes strong selection on the alphavirus nsP3/4 opal codon. **(A)** Alphaviruses encode a premature in-frame opal codon within the non-structural open reading frame at the 3’ end of the nsP3 gene. **(B)** Infection rates of opal-to-sense substitution SINV variants relative to wild-type SINV in Aedes albopictus cells: either U4.4 cells with functional Dcr2 (*Dcr2+*) or CRISPR/Cas9-edited Dcr2-deficient (*Dcr2 KO*) U4.4 cells, or (*Dcr2-*) C6/36 cells. Data represent the mean of at least three independent biological replicates. **(C)** Competitive fitness of SINV 550C variant against wild-type SINV 550opal in *Dcr2+* U4.4 or *Dcr2-* C6/36 cells. Wild-type versus wild-type competition was performed as a control. The data represent four independent biological replicates. Error bars represent the standard error of the mean (SEM). Two-way ANOVA with Tukey’s test for multiple comparisons. **** = p < 0.0001, *** = p < 0.001, ** = p < 0.01, * = p < 0.05, ns = not significant. **(D)** Wild-type SINV (with opal codons, UGA) or 550C SINV variants (with UGC/UGU codons) were passaged for 5 days in *Dcr2+* U4.4 cells or CRISPR-edited U4.4 *Dcr2 KO* cells. The frequencies of opal versus cysteine codons at the start and end of the passaging, as determined by small RNA sequencing, are reported. The data represent three independent biological replicates.

Considerable interest has focused on how and why the opal codon confers a fitness advantage to alphaviruses (*4–7*). We recently demonstrated that the opal codon provides a fitness advantage in vertebrate cells by resolving a temperature-dependent trade-off between nsP4 expression via PRT and the efficiency of viral nsP (*1*). We showed that the opal codon at nsP3 position 550 (550opal) is strongly preferred over non-opal (sense codon) substitutions at this site in vertebrate cells grown at 37°C. However, why this opal codon is maintained in insect hosts remains unknown, since, in contrast to vertebrate cells at 37°C, non-opal substitutions are much better tolerated in vertebrate cells or *Aedes albopictus*-derived C6/36 mosquito cells grown at 28°C (*1*, *7*, *8*). Moreover, previous studies with distinct alphaviruses, including SINV and Eastern equine encephalitis virus (EEEV), have shown that Opal-to-Cysteine substitutions may confer a replicative fitness advantage over the wild-type opal codon in mosquito C6/36 cells (*1*, *9*). Despite this apparent fitness benefit, opal-to-sense codon substitutions are exceedingly rare (3%) in alphaviruses isolated from mosquitoes and are entirely absent in closely related insect-specific alphaviruses (Figure S1A-B) (*10–12*). Thus, additional host or environmental factors must contribute to the conservation of the nsP3 opal stop codon in insect hosts.

A key factor that could select for preservation of the opal codon among alphaviruses is innate immunity in insect hosts. Alphaviruses elicit a robust small RNA response in mosquito cells, mediated primarily by the siRNA pathway (*13–15*). Following viral infection, Dicer 2 (Dcr2), a key mediator in the short interfering RNA (siRNA) pathway, processes viral double-stranded RNA (dsRNA) replication intermediates into 21-nt virus-derived small interfering RNAs (vsiRNAs). These vsiRNAs assemble with Argonaute 2 (Ago2) and other components of the RNA-induced silencing complex (RISC) to target nascent single-stranded viral RNA for cleavage and degradation (*16*, *17*). However, most previous experimental passaging studies were conducted in C6/36 cells, which lack an effective antiviral RNA interference (RNAi) response due to impaired Dcr2 function (*18*).

We investigated whether immune pressure from RNAi might explain the retention of the opal codon in insect hosts, using the prototype alphavirus Sindbis virus (SINV) as our model and genetic perturbations of RNAi pathway components in mosquito cells. Our previous study identified a polyprotein processing defect associated with viruses carrying opal-to-sense substitutions. Here, we asked whether such processing defects also alter the integrity of viral replication spherules, thereby allowing host cytoplasmic RNA nucleases (such as Dicer in insect cells) and sensors (such as RIG-I-like proteins in vertebrate cells) to detect viral RNA and trigger cellular immune responses. To test this, we used cytological assays to quantify viral spherule integrity and to assess host antiviral responses, including siRNA production in insect cells and interferon (IFN) production in vertebrate cells. Our study demonstrates that by ensuring proper polyprotein processing, the conserved nsP3 opal stop codon protects alphaviruses from intracellular immune defenses across evolutionarily divergent hosts.

## Results

### The Dcr2-dependent siRNA pathway imposes selection against non-opal codon variants

*Aedes albopictus*-derived C6/36 cells lack functional Dicer 2 (Dcr2) due to a frameshift mutation in the Dcr2 open reading frame, resulting in a Dcr2 protein that lacks the RNase III domains essential for dicing activity (*18*). Prior work from our lab and others has shown that alphaviruses can tolerate certain opal-to-sense codon substitutions in mosquito cells lacking a functional Dcr2-dependent siRNA pathway which is the primary antiviral response against alphavirus infection in mosquitoes (*1*, *8*, *9*). To determine whether Dcr2 deficiency accounts for tolerance of opal codon substitutions in C6/36 cells, we tested the growth of three opal-to-sense substitution variants (550Alanine, 550Arginine, 550Cysteine) relative to wild-type SINV (550opal) in Dcr2-competent U4.4 cells or Dcr2-lacking C6/36 cells. To ensure quantifiable virus growth in RNAi-competent cells, all infections were performed at a moderately high MOI (≥4). Consistent with our previous study (*1*), we found that infection rates of the 550C and 550A variants were higher than those of wild-type SINV (550opal) in Dcr2-lacking C6/36 cells, whereas the SINV 550R variant was less fit. In contrast, infection rates of all sense-codon variants were significantly reduced in Dcr2-competent U4.4 cells compared to wild-type SINV (550opal) (Figure 1B).

Both U4.4 and C6/36 cells are derived from *Aedes albopictus*. Nevertheless, differences among these cell lines, beyond their Dcr2 status, could influence SINV infection. To compare the fitness of viral variants within isogenic host cell lines, we used CRISPR/Cas9 to generate *Dcr2*-knockout (*Dcr2 KO*) U4.4 cells, yielding a heterogeneous population in which *Dcr2* was knocked out in ∼82-85% of cells (Figure S2). Despite the incomplete nature of this knockout, infection rates of all SINV opal-to-sense variants, including SINV 550R, were comparable (550A, 550R) or higher (550C) than wild-type SINV (550opal) (Figure 1B). Thus, in both C6/36 and U4.4 cells, loss of *Dcr2* enhances the fitness of SINV opal-to-sense variants.

Our recent detailed analysis of the SINV opal codon in *Dcr2*-deficient mosquito cells revealed cysteine as the most tolerated sense codon (*1*). Cysteine is also the only opal-to-sense substitution observed in SINV in nature (*19*). Therefore, we conducted all subsequent comparative analyses using wild-type SINV (550opal) and SINV 550C. Consistent with the independent growth assays, our competition experiments demonstrated that SINV 550C has significantly lower fitness than 550opal in *Dcr2*-competent U4.4 cells, but higher fitness than wild-type SINV in *Dcr2*-lacking C6/36 cells (Figure 1C), confirming our previous findings (*1*).

We also assessed the selective pressure exerted by intact Dcr2 by tracking SINV 550C variants across three independent replicate lineages over a five-day growth period, which roughly corresponds to 24 rounds of replication (Figure 1D). We observed that SINV 550C was nearly completely replaced by 550opal in *Dcr2*-competent U4.4 cells, indicating that mutations reverting 550C to 550opal had occurred and spread through the population within just 24 rounds of replication. In contrast, 550C variants persisted in U4.4 *Dcr2* KO cells (Figure 1D). Conversely, WT SINV 550opal almost entirely converted to 550sense variants, primarily 550C (UGC or UGU codons), within 5 days in U4.4 *Dcr2* KO cells. However, WT SINV 550opal variants persisted in *Dcr2*-competent U4.4 cells (Figure 1D). The swift accumulation of opal-to-sense substitutions in *Dcr2* KO U4.4 cells mirrors previous findings in Eastern equine encephalitis virus (EEEV), which also develops opal-to-Cys mutations after long-term passaging in *Dcr2*-deficient C6/36 cells (*20*). Overall, our results show that Aedes Dcr2 applies strong selective pressure to maintain the SINV opal codon (Figure 1).

We next tested whether the *in vitro* fitness differences between SINV 550opal and SINV 550C translated into altered infection outcomes in *Aedes* mosquitoes. For this, we used two reagents. The first is a previously published *Aedes aegypti Dcr2* loss-of-function mutant generated by TALEN, in which a constitutively expressed eGFP knock-in cassette disrupts *Dcr2* (*21*). Second, we generated a CRISPR/Cas9-mediated *Dcr2* knockout line with a constitutively expressed *dsRED* inserted into *Dcr2*, thereby disrupting the open reading frame (Figure 2A) (*22*). Trans-heterozygous expression of eGFP and dsRED allows us to visually track the independent insertions in loss-of-function mutants. By crossing the independent heterozygous *A. aegypti* lines, we generated a trans-heterozygous *Dcr2* knockout (yellow, Figure 2B). We tested *Dcr2* loss of function by analyzing survival curves following systemic infection with wild-type SINV (550opal). Consistent with previous data in homozygous *Dcr2-*null eGFP mosquitoes, trans-heterozygous *Dcr2-*null mosquitoes showed a lethal phenotype compared to their wild-type siblings (Figure 2C) (*21*, *22*).

**Figure 2.**
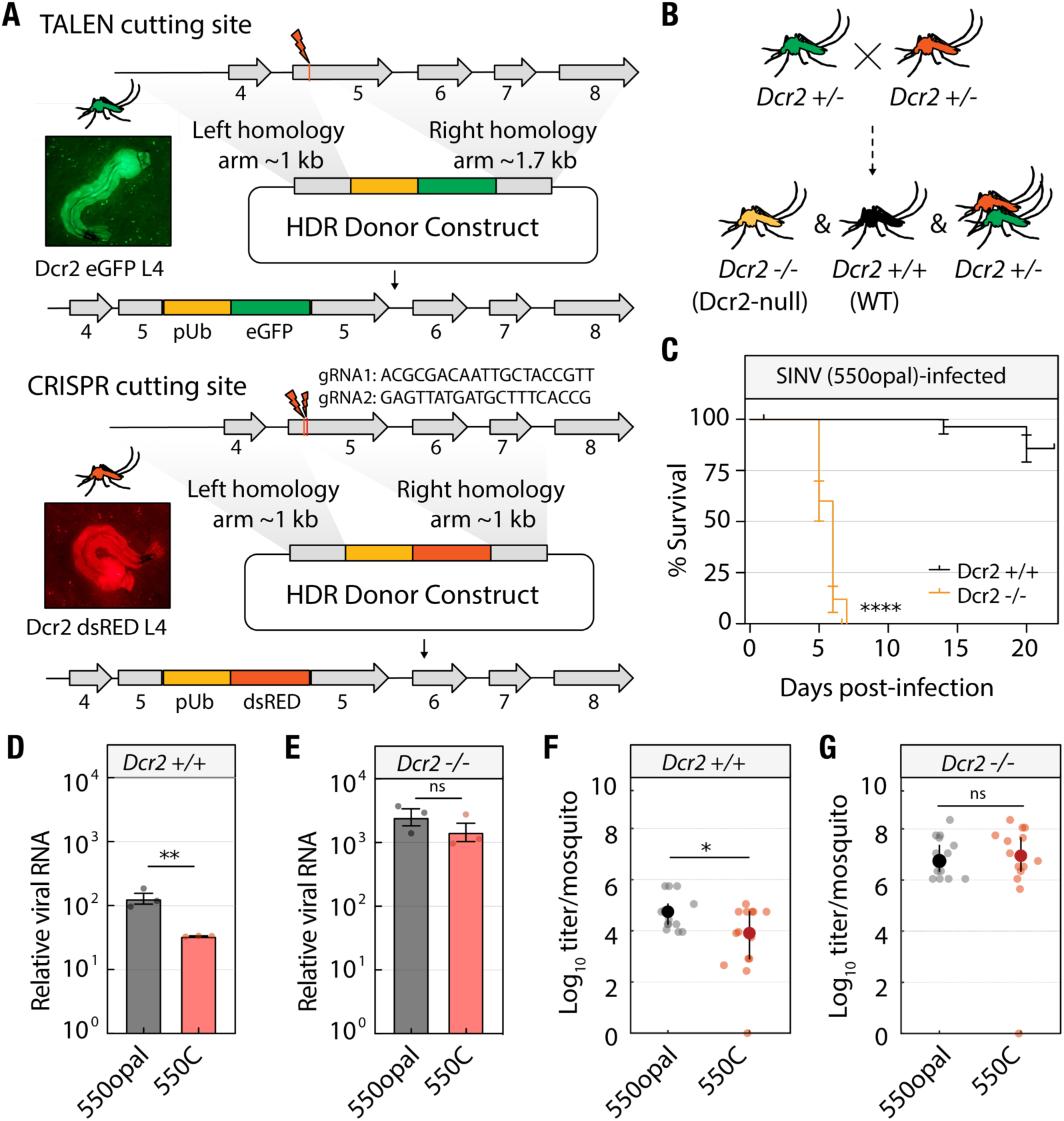
The mosquito RNAi pathway imposes strong selection on SINV nsP3 opal codon *in vivo*. **(A-B)** (Top) Schematic of TALEN-mediated HDR integration to generate *Dcr2*-null transgenic lines with the knock-in of a frame-disrupting pUb-eGFP marker. (Middle) Schematic of CRISPR-Cas9-mediated *Dcr2*-null transgenic lines with the knock-in of a frame-disrupting pUb-dsRED marker. Photographs are representative of the eGFP, or dsRED-expressing mosquitoes selected for crossing experiments **(B)** to generate wild-type (*Dcr2*+/+, black) or trans-heterozygous *Dcr2*-null (*Dcr2*−/−, yellow) mosquitoes (Bottom). **(C)** Survival of sibling wild-type (black line) or trans-heterozygous *Dcr2-*null (yellow line) mosquitoes following systemic SINV infections. Survival curves represent cohorts of ≥28 adult female mosquitoes infected with 10^6^ TCID_50_/ml of wild-type SINV (550opal). Significance was determined using the log-rank (Mantel-Cox) test (p-value < 0.0001). **(D-E)** Wild-type (*Dcr2*+/+) or *Dcr2*-null (*Dcr2*−/−) *Aedes aegypti* mosquitoes were infected with SINV 550opal or 550C. Three days post-infection, viral RNA was quantified via qRT-PCR. The data represent three independent biological replicates, each consisting of five pooled mosquitoes. Reported values are normalized to 550opal RNA levels in wild-type mosquitoes. **(F-G)** Viral loads were assessed from individual **(D)** wild-type (*Dcr2*+/+, black) or **(E)** homozygous *Dcr2*-null (*Dcr2*−/−, yellow) mosquitoes (n=15) collected three days post-infection. Error bars represent the standard error of the mean (SEM). Unpaired t-tests. ** = p < 0.01, * = p < 0.05, ns = not significant.

We then challenged wild-type or *Dcr2−/−* knockout sibling *A. aegypti* mosquitoes with equal doses of wild-type SINV (550opal) or the 550C SINV variant (Figure S3). Consistent with our *in vitro* findings, total viral RNA levels in SINV 550C-infected wild-type mosquitoes were 4-fold lower than those in SINV 550opal-infected wild-type mosquitoes three days post-infection (Figure 2D). This indicates that SINV 550C experiences a significant fitness loss in wild-type mosquitoes. As expected, loss of *Dcr2* significantly increased infection levels for both wild-type SINV (550opal) and SINV 550C. Importantly, viral RNA levels in SINV 550C-infected *Dcr2*−/− mosquitoes were statistically indistinguishable from those in wild-type SINV (550opal)-infected *Dcr2*−/− mosquitoes (Figure 2E).

We also assessed *in vivo* viral fitness by quantifying the infectious virus titer recovered from infected mosquitoes. Consistent with total viral RNA levels, we recovered five-fold less infectious virus from SINV 550C-infected *Dcr2+/+* mosquitoes than from those infected with wild-type SINV (550opal) (Figure 2F). As expected, we recovered significantly more infectious virus from *Dcr2−/−* mosquitoes for both viruses (Two-way ANOVA with Tukey’s multiple-comparison test (p < 0.01 for 550opal; p < 0.0001 for 550C). In addition, the levels of recovered virus were equivalent between wild-type SINV (550opal) and SINV 550C-infected *Dcr2−/−* mosquitoes (Figure 2G). These findings confirm that the SINV opal codon confers a significant replicative advantage *in vivo* and that RNAi loss rescues the *in vivo* fitness defects of the SINV 550C variant (*21*, *22*). Our present findings in SINV echo previous findings on ONNV in *Anopheles gambiae* mosquitoes (*4*).

### Opal-to-Cys substitution induces higher small RNA response in Dcr2-competent mosquito cells

Because SINV 550C exhibits reduced fitness in *Dcr2*+ cells, we hypothesized that this variant might elicit a more pronounced siRNA response than wild-type SINV (550opal). To test this, we performed small RNA sequencing of *Dcr2*+ U4.4 or *Dcr2*– C6/36 cells infected with either 550opal or 550C SINV variants (Figure 3A). *Dcr2 KO* U4.4 cells were excluded from the small RNA sequencing analysis because the knockout was incomplete. Although SINV 550C infected nearly 10 times fewer cells than wild-type SINV (Figure 3A), virus-derived small RNA levels were significantly higher in *Dcr2*+ cells infected with SINV 550C than in those infected with wild-type SINV (550opal) (Figure 3B). This observation directly links the lower growth of SINV 550C in *Dcr2*+ cells to an elevated immune response. Therefore, a single nucleotide change (UGA to UGC) in the 11.7-kb SINV genome, which causes the loss of the nsP3 opal codon, is enough to produce a significantly higher small RNA response in mosquito cells.

**Figure 3.**
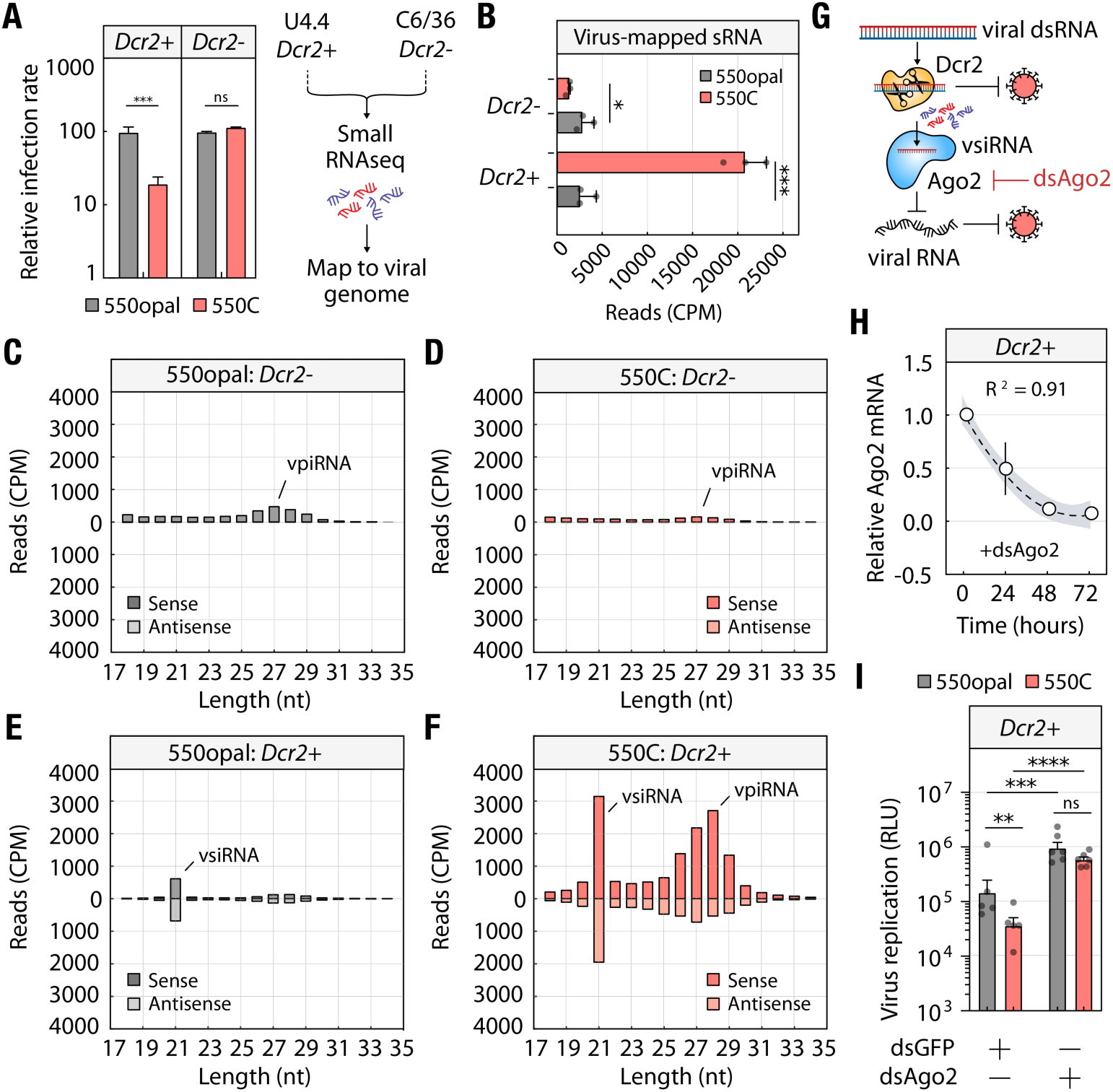
SINV sense-codon variant induces higher small RNA response in mosquito cells. **(A)** *Dcr2*+ U4.4 or *Dcr2*– C6/36 cells were infected with wild-type SINV 550opal or the SINV 550C variant. Relative infection rates of SINV 550opal and 550C in *Dcr2+* and *Dcr2-* cells used for Small RNAseq. Small RNA sequencing was performed on cells collected five days post-infection. **(B)** Normalized small RNA read counts mapped to SINV genomes in *Dcr2*+ U4.4 or *Dcr2*– C6/36 cells infected with wild-type SINV 550opal and SINV 550C (n=3). **(C-D)** Size distribution of small RNAs derived from *Dcr2*- C6/36 cells infected with wild-type SINV 550opal or SINV 550C. **(E-F)** Size distribution of small RNAs derived from *Dcr2*+ U4.4 cells infected with wild-type SINV 550opal or SINV 550C. The data represent three independent biological replicates. **(G)** Schematic of the mosquito RNAi pathway and the roles of *Dcr2* and *Ago2* in virus restriction. **(H)** Quantification of relative *Ago2* transcript levels by qRT-PCR in *Dcr2*+ mosquito cells (n=3) treated with *Ago2* double-stranded RNA (dsAgo2). The dotted line denotes the expected trend from a one-phase exponential decay model. Bands represent the 95% confidence interval. **(I)** Effect of dsAgo2 treatment on the replication of wild-type SINV 550opal and SINV 550C in *Dcr2*+ mosquito cells as quantified by luciferase assay. Error bars represent the standard error of the mean (SEM). The data represent five independent biological replicates. Student’s t-tests. *** = p < 0.001, * = p < 0.05, ns = not significant.

To further analyze the nature of the small RNA response, we examined small RNAs by size distribution and polarity (Figure 3C-F, S4A-D). As expected, the siRNA response to either SINV 550C or SINV 550opal was significantly reduced in C6/36 cells lacking *Dcr2* (Figures 3C-D). Additionally, all small RNAs in C6/36 cells showed a bias toward the sense strand. In Dcr2-competent U4.4 cells, levels of 21-nucleotide vsiRNAs were notably higher in SINV 550C-infected cells compared to 550opal-infected cells (Figures 3E-F), consistent with SINV 550C’s increased susceptibility to Dcr2-mediated processing (Figures 3B). We also detected clear differences in siRNA polarity between the two viral variants. siRNAs originating from 550opal mapped equally to sense and antisense viral RNA, indicating they arise from double-stranded RNA replication intermediates, where sense and antisense RNAs are in equal amounts (paired t-test: p = 0.976; Figure 3E). Conversely, 550C-derived siRNAs showed a strong bias toward the sense strand (paired t-test: p < 0.05, Figure 3F). Our results suggest that siRNAs produced in SINV 550C-infected cells are more plentiful than those in wild-type SINV (550opal) infected cells and stem from single-stranded positive-sense viral RNA rather than double-stranded RNA. We considered two complementary mechanisms by which increased siRNA production during SINV 550C infection in *Dcr2*-competent U4.4 cells could directly reduce fitness (Figure 3G, top). First, Dcr2-mediated cleavage of viral dsRNA could inherently harm the virus by disrupting minus-strand replication. This could lead to a proportional reduction in intact viral dsRNA, which might be enough to lower viral fitness. Alternatively, viral siRNAs produced by Dcr2 can be targeted to degrade new viral RNA via Argonaute 2 (Ago2), thereby restricting viral replication (Figure 3G, bottom). To distinguish between these two mechanisms, we tested whether Ago2-mediated targeting of vsiRNAs to viral RNAs is necessary for SINV 550C inhibition. We performed *Ago2* knockdown in *Dcr2*+ U4.4 cells using double-stranded RNA (dsRNA), achieving over 90% knockdown within 48 hours (Figure 3G-H, Figure S5A). If both cleavage and Ago2 targeting play significant antiviral roles, then Ago2 knockdown should only modestly affect viral fitness (Figure 3G). However, contrary to this expectation, Ago2 knockdown greatly increased SINV 550C replication to levels similar to wild-type SINV (550opal) (Figure 3I). This shows that Ago2-mediated vsiRNA targeting of viral RNA significantly contributes to the reduced fitness of SINV 550C in *Dcr2*+ cells.

In addition to vsiRNAs, we also detected viral-derived 24-31 nt piwi-interacting RNAs (piRNAs) in both C6/36 and U4.4 cells, consistent with previous studies (Figures 3C-F, S6, S7, S8A) (*14*, *15*, *22*). In mosquito cells, viral piRNAs (vpiRNAs) are primarily generated via a ping-pong-like mechanism involving specific PIWI proteins, including Ago3 and Piwi5 (*23–25*). Consistent with earlier findings, we observed that SINV-derived piRNA reads primarily originate from the subgenomic RNA, especially from a hotspot at the 5’ end of the capsid gene (Figure S6B), through a canonical ping-pong amplification pathway, as indicated by the 1U bias in antisense and 10A bias in sense piRNAs (Figure S8C). Like siRNAs, C6/36 cells produce significantly fewer viral-derived piRNAs than U4.4 cells (Figures 3C-D, S6). *Dcr2*+ U4.4 cells infected with the SINV 550C variant also produce 24-31 nt viral-derived piRNAs at levels 12 times higher than the wild-type SINV variant (Figures 3E-F and S8A). Conversely, SINV 550C infection leads to a 2-fold decrease in small RNAs in mosquito cells lacking Dcr2, which show an almost complete absence of vsiRNA and vpiRNA production (Figure 3B-C). Therefore, although vpiRNA production is traditionally considered independent of mosquito Dcr2 activity, our results add to the growing evidence of a functional interaction between the siRNA pathway and ping-pong-generated piRNA production in mosquitoes (*24*, *26–28*). Unlike viral-derived siRNAs, which have been shown to restrict alphavirus replication in insect cells, the antiviral role of viral-derived piRNAs remains less clear (*29*). In our study, we observed no significant differences in the levels of subgenomic RNA template or structural protein expression between wild-type SINV (550opal)- infected and SINV 550C-infected *Dcr2*+ cells (Figure S8D). Moreover, since knockdown of *Ago2*, which is involved in siRNA but not piRNA targeting, was sufficient to restore the fitness of the SINV 550C variant to wild-type levels in U4.4 cells (Figure 3I), we conclude that viral-derived piRNAs do not play a substantial antiviral role and do not discuss them further in this study.

### SINV Structured RNA element protects against RNAi in mosquito cells

To determine whether specific SINV genomic regions serve as Dcr2 substrates in wild-type SINV and, especially in cells infected with SINV 550C, we mapped all 21-nt vsiRNA reads to the SINV reference genome. Using a 40-nucleotide sliding window across the SINV genome, we identified regions with higher-than-average vsiRNA coverage (Figure 4A). Since infection with the SINV 550C variant induces a stronger siRNA response than wild-type SINV (550opal), we mainly focused on regions with significantly higher vsiRNA coverage in SINV 550C compared to wild-type SINV (550opal) (Figure 4B). This differential analysis uncovered several windows in the SINV genome that produced disproportionately more vsiRNAs during infection with the SINV 550C variant. The most notable was a 23-nt stretch within the E1 structural gene coding region. This ‘hotspot,’ which we named E1-hs, made up more than 4% of total siRNA reads in SINV 550C (99.85th percentile). E1-hs is also a hotspot in SINV 550opal, contributing 1% of total siRNA reads (96th percentile) (Figures 4A-B).

**Figure 4.**
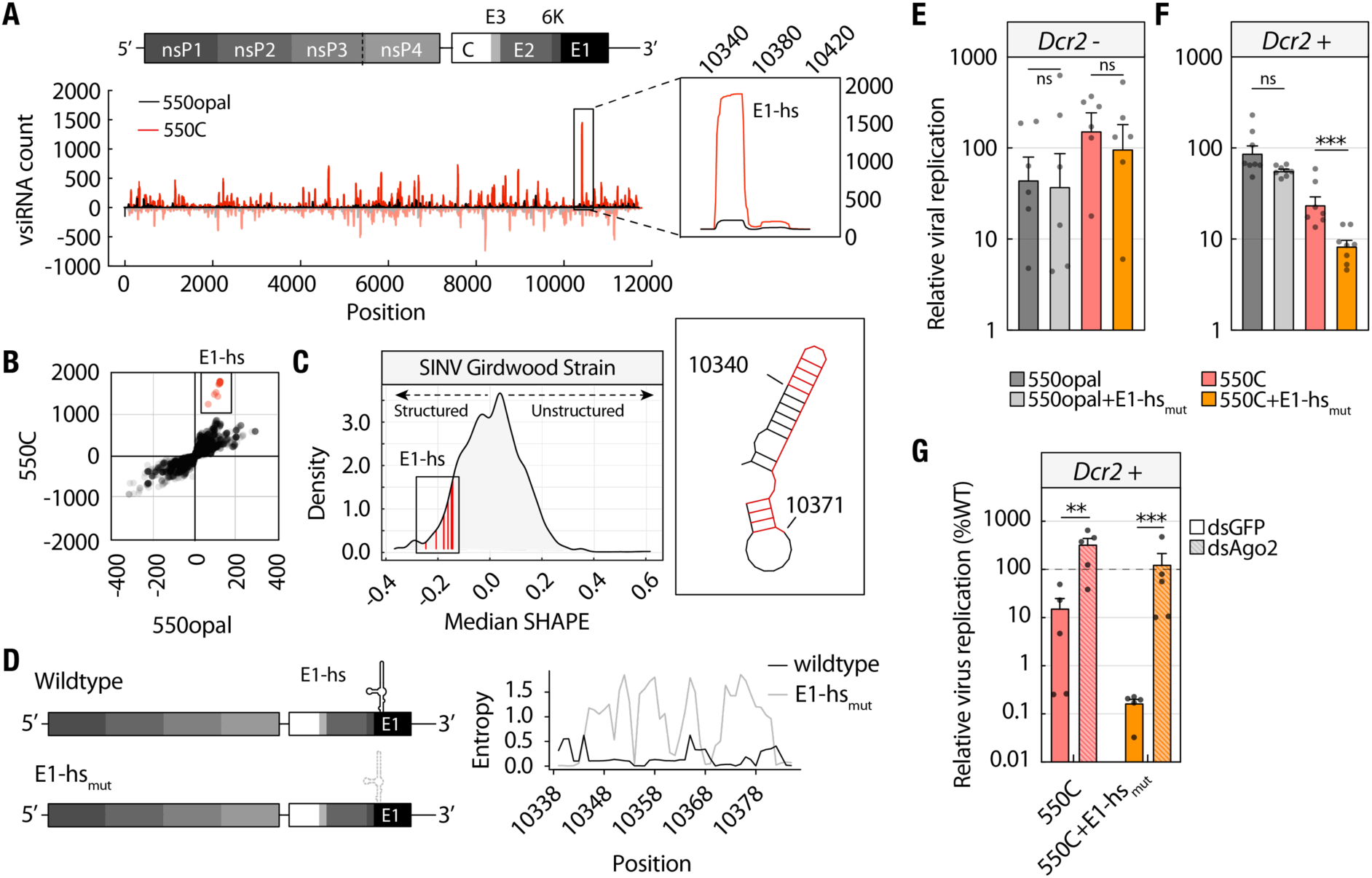
A structured RNA element enables SINV to blunt the siRNA response. **(A)** Distribution of vsiRNA reads from wildtype SINV 550opal (in black) or SINV 550C (in red) viruses in *Dcr2*+ U4.4 cells. Positive and negative Y-axis values represent read counts mapped to the sense and antisense strands, respectively, at every position along the SINV genome (X-axis). The inset shows a zoomed-in view of the E1-hs region. **(B)** Correlation of per-site vsiRNA reads mapped to wildtype SINV 550opal or SINV 550C viruses. Residues within the E1-hs region are boxed and colored in red. **(C)** Density distribution profile of median SHAPE values of SINV RNA. Median SHAPE values of residues within the E1-hs region are boxed and highlighted in red. Location of the E1-hs region in SHAPE-constrained RNA secondary structure. **(D)** Schematic of viral genomes with intact (wildtype) and mutated E1-hs (E1-hs_mut_). Positional entropy profile of E1-hs RNA structural element in wild-type SINV and the E1-hs_mut_ strain. High and low entropy values indicate conformationally flexible and rigid structures, respectively. **(E)** Replication of wildtype SINV 550opal or SINV 550C viruses with intact or destabilized E1-hs structure in *Dcr2*– C6/36 cells as quantified via luciferase reporter assays. The data represent six independent biological replicates. **(F)** Replication of wildtype SINV 550opal or SINV 550C viruses with intact or destabilized E1-hs structure in *Dcr2*+ U4.4 cells. The data represent six independent biological replicates. **(G)** Relative replication levels of SINV 550C viruses with intact or destabilized E1-hs structure in *Dcr2*+ U4.4 cells in the presence or absence of Ago2, normalized to wild-type SINV replication (550opal; horizontal dotted line). The data represent values from five independent biological replicates. Two-way ANOVA with Tukey’s multiple comparisons test. **** = p < 0.0001, ** = p < 0.01, * = p < 0.05, ns = not significant. Two-way ANOVA with Tukey’s multiple comparisons test. **** = p < 0.0001, *** = p < 0.001, * = p < 0.05, ns = not significant.

The SINV RNA genome contains several structural RNA elements vital for replication, expression, and packaging (*30*). Due to the biased polarity of vsiRNAs derived from SINV 550C, these elements could serve as hotspots for disproportionately large amounts of viral-derived siRNAs. In fact, vsiRNA reads mapping to the E1-hs region showed a strong bias toward the sense strand (Figure 4A inset), suggesting that these E1-hs siRNAs may originate from a structured RNA element in the SINV genome. To determine whether E1-hs corresponds to a structured RNA, we analyzed previously generated SINV RNA SHAPE-MaP data by Kutchko et al. to calculate and visualize the median SHAPE reactivity values across a 40-nucleotide sliding window throughout the SINV genome (*30*). Residues within the E1-hs region displayed notably negative SHAPE reactivity values, indicating they are part of highly structured RNA in SINV (Figure 4C). We determined the RNA secondary-structure conservation of the E1-hs element using RNAalifold and a multiple-sequence alignment of 2,086 alphavirus sequences collected from BV-BRC. We then mapped the corresponding E1-hs region to the CHIKV SHAPE MaP data (Figure S9C) described in Madden et al (*31*). Thus, as with SINV, the E1-hs sequence also appears to be part of structured RNA in CHIKV (Figure S9C) and likely in other alphaviruses. Additionally, we compared genome-wide vsiRNA profiles between SINV 550C and SFV using previously collected data from Siu et al. from SFV4-infected *Dcr2*+ U4.4 *(Aedes albopictus)* and Aag2 *(Aedes aegypti)* cells (*32*). This comparison identified several hotspots of vsiRNA production, including the SINV E1-hs region, which is shared by both viruses despite their high overall sequence divergence (Figure S10A-B).

To evaluate the functional importance of the E1-hs hotspot, we introduced synonymous mutations that disrupt the E1-hs secondary structure without changing the E1 protein-coding sequence (Figure 4D). We then compared the replication efficiency of these E1-hs_mut_ variants with that of their wild-type counterparts in both SINV 550C and SINV 550opal viruses in *Dcr2*– (C6/36) cells. In *Dcr2*–cells, SINV E1-hs_mut_ replicated at levels indistinguishable from wild-type SINV, and SINV 550C E1-hs_mut_ replicated at levels comparable to SINV 550C (Figure 4E). Thus, E1-hs_mut_ variants do not intrinsically impair viral replication. We then investigated the effect of the E1-hs mutation in *Dcr2*+ cells. Although the replication of the SINV 550opal E1-hs_mut_ variant was comparable to that of SINV 550opal, SINV 550C E1-hs_mut_ replicated to significantly lower levels than SINV 550C in *Dcr2*+ cells (Figure 4F). Replicative fitness of SINV 550C E1-hs_mut_ was also rescued to wild-type SINV levels in *Dcr2*+ cells following *Ago2* knockdown, indicating that E1-hs helps sustain viral fitness under active RNAi targeting (Figure 4G). Thus, the ability of E1-hs to produce excess vsiRNAs is vital for preserving the fitness of the 550C variant in *Dcr2*+ cells, since the SINV 550C genome is more vulnerable to Dcr2 targeting. Our results show that the SINV opal codon and the E1-hs structure work together as a powerful RNAi evasion strategy.

### Disrupted nsP processing compromises the integrity of viral replication spherules and increases susceptibility to mosquito RNAi

We next asked why the SINV 550C variant is much more susceptible to RNAi than wild-type SINV (550opal). We previously discovered that opal-to-sense substitutions cause overproduction of the full-length nsP (P1234), thereby disrupting polyprotein processing kinetics. These differences in polyprotein processing between wild-type SINV and sense-codon variants are most prominent in vertebrate cells cultured at 37°C but also appear (though less strongly) in both vertebrate and mosquito cells maintained at 28°C (*1*). Specifically, less P123 is produced early during infection in SINV 550C-infected *Dcr2*+ cells, presumably due to delayed P3/4 processing caused by excess P1234 (Figure S11A-B). We therefore considered whether functional RNAi could worsen delays in polyprotein processing. However, similar polyprotein processing trends were seen in *Dcr2*+ (U4.4) and *Dcr2*– (C6/36) mosquito cells at 28°C (Figure S11A and C, S11B and D), ruling out the possibility that RNAi status influenced SINV nsP polyprotein processing.

The SINV nsP1 I538T mutation also slows P1/2 processing, and we recently showed that it helps restore the disruption in nsP processing cadence caused by the 550C mutation by creating a ‘double delay’ (*1*). Therefore, we tested whether SINV 550C replication is improved by the nsP1 I538T mutation in *Dcr2*+ cells. However, we did not observe any appreciable improvement in the replication of SINV 550C+I538T compared to SINV 550C, suggesting that the ‘double delay’ may not enhance spherule integrity defects in a timely manner to prevent RNAi susceptibility (Figure S11E).

Next, we investigated whether the increased RNAi susceptibility of the SINV 550C variant was due to increased viral dsRNA production, which could lead to higher antiviral vsiRNA output in *Dcr2*+ cells. This increase in dsRNA could directly result from the nsP processing defect, which delays transitions from minus-strand to plus-strand replication. We have previously used a quantitative RT-PCR (qRT-PCR) assay to measure intracellular levels of SINV genomic RNA, including both plus and minus strands (*1*). However, we were concerned that Dcr2 cleavage of viral dsRNA might prevent accurate measurement of viral dsRNA levels within cells. Therefore, to detect both intact and potentially processed dsRNA, we used the J2 monoclonal antibody, which has been well characterized for detecting dsRNAs ranging from 14 to 40 bp (*33*). We performed immunofluorescence microscopy on *Dcr2*+ (U4.4) cells, which revealed distinct J2 foci in the cytoplasm of SINV-infected cells but not in uninfected cells. These J2 foci indicate SINV replication compartments (RCs) or spherules (Figure 5A). We measured J2/dsRNA intensity within these spherules. Although the total number of spherules per cell did not change, *Dcr2*+ (U4.4) cells infected with SINV 550C showed a 29% increase in spherule J2 intensity compared to those infected with wild-type SINV (550opal), suggesting that SINV 550C replication spherules contain more J2-binding dsRNA substrate (Figure 5B-C).

**Figure 5.**
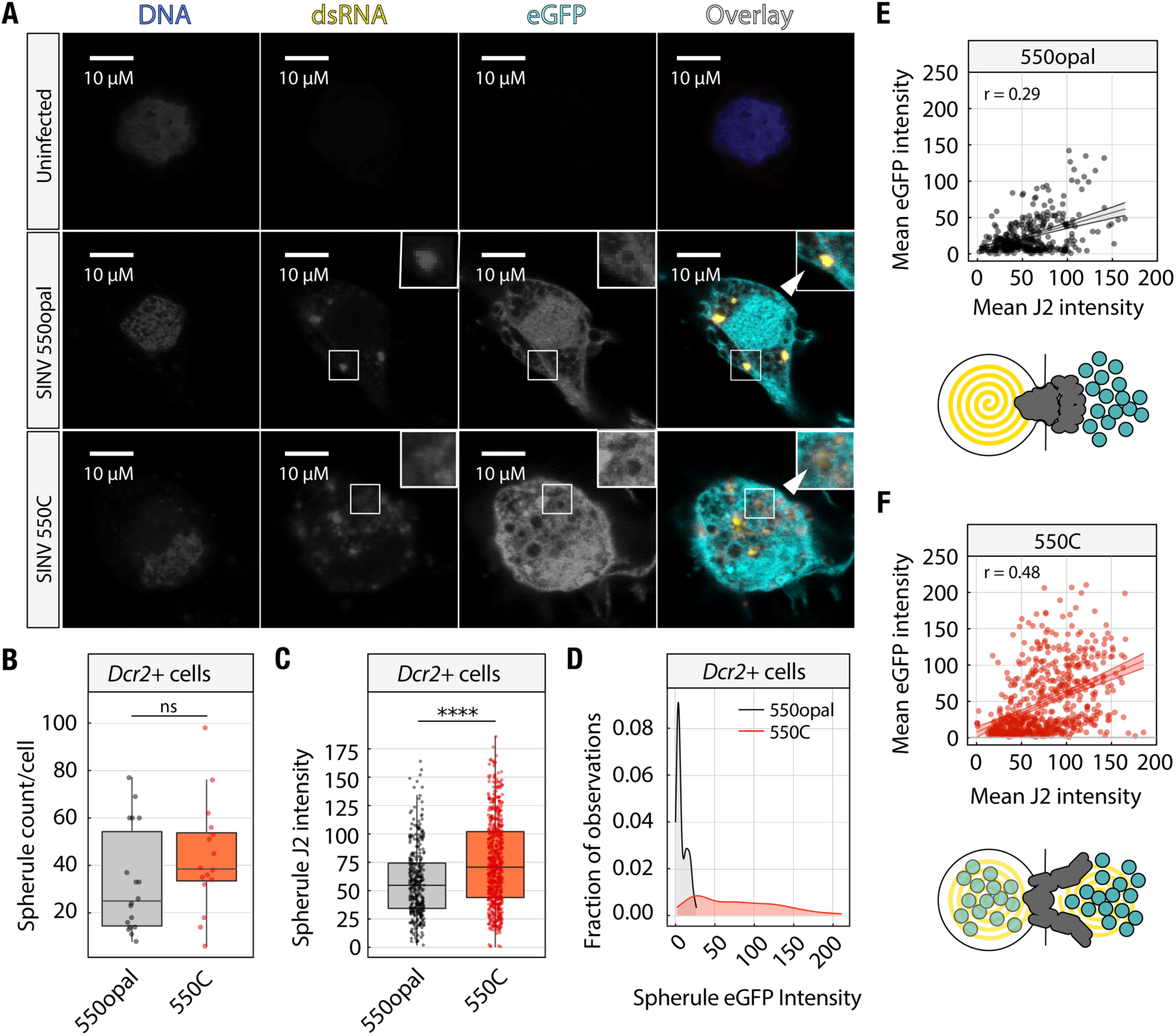
Delayed non-structural polyprotein processing in SINV 550C compromises the integrity of viral replication spherules. **(A)** Confocal microscopy-based localization of viral dsRNA (J2) and virally encoded eGFP in *Dcr2*+ U4.4 cells that were either uninfected (top row), infected with wild-type SINV 550opal (middle row), or infected with SINV 550C (bottom row). The inset shows the localization of eGFP and dsRNA around (middle row) or within (bottom row) viral replication spherules. White arrows indicate spherule outer boundaries. **(B)** Spherule count per cell (n=17) is reported as the mean number of spherules per cell in each z-stack. **(C)** Quantification of J2 intensities within viral replication spherules (n=629) in *Dcr2*+ U4.4 cells infected with either wild-type SINV 550opal or SINV 550C. Mann-Whitney U test. **** = p < 0.0001, ns = not significant. Error bars represent the standard error of the mean (SEM). **(D)** Density distributions of observed values of spherule eGFP intensity in wild-type SINV 550opal or SINV 550C-infected *Dcr2*+ cells. Kolmogorov-Smirnov (KS) test = 0.862, p < 0.0001. **(E-F)** Correlation between mean dsRNA and eGFP intensities per spherule across *Dcr2*+ cells infected with either wild-type SINV 550opal or SINV 550C. Spearman’s correlation tests (E-F: p < 0.0001).

Recent cryo-EM reconstruction of wild-type alphavirus replication organelles showed that each spherule contains only a single full-length viral dsRNA molecule (*34*). We therefore wondered whether the accumulation of Dcr2-processed dsRNA fragments could increase spherule J2 intensity in SINV 550C-infected *Dcr2*+ cells. To test this, we performed a similar analysis in *Dcr2*- C6/36 cells (Figure S12A). We reasoned that if SINV 550C naturally synthesizes excess full-length dsRNA, we would also expect a similar increase in spherule J2 signal in cells lacking Dcr2. However, we found no significant differences in spherule J2 intensities between wild-type and SINV 550C-infected C6/36 cells (Figure S12B-C). These findings confirm both our previous small RNAseq analyses showing greater siRNA production in SINV 550C-infected U4.4 cells and our hypothesis that the increased spherule J2 signal in SINV 550C-infected U4.4 cells (Figure 5B-C) reflects an excess of processed dsRNA fragments.

Another consequence of nsP processing defects in cells infected with the SINV 550C variant could be compromised integrity or permeability of mature viral replication spherules, which are composed of processed non-structural proteins. Initial alphavirus replication spherule formation requires the intact P123 polyprotein, while spherule maturation depends on further processing into nsPs 1-4 (*35*, *36*). Formation of replication spherules is a passive but highly effective viral evasion tactic that hides viral dsRNA replication intermediates from cellular RNA sensors and effectors, including Dcr2 (*37*, *38*). The presence of processed dsRNA within or near replication spherules in SINV 550C-infected *Dcr2*+ cells suggests that mature spherule formation is delayed in SINV 550C because of abnormal processing of the P123 polyprotein, which weakens the structural integrity of replication complexes, increases spherule permeability, and allows cytoplasmic Dcr2 access to cleave viral dsRNA. To test this idea, we measured spherule integrity by assessing their ability to block cytoplasmic eGFP.

If viral spherule integrity were intact, we would expect to see occlusion of the normally freely diffusing cytoplasmic eGFP. Indeed, 2D plot profile analysis of dsRNA-containing intact spherules (identified using the J2 antibody) in *Dcr2*+ cells infected with eGFP-expressing wild-type SINV (550opal) showed a clear reduction in eGFP intensity within spherules compared to areas outside the spherule boundary, indicating the selective exclusion of cytoplasmic eGFP from the viral spherule compartment (Figure S12D). In contrast, within-spherule eGFP intensity was markedly higher in SINV 550C-infected cells. When we examined a larger number of J2-marked viral replication spherules in *Dcr2*+ cells infected with wild-type SINV (550opal), we observed a bimodal distribution of spherule eGFP intensity values: most spherules exclude eGFP entirely, while only a very small subgroup of spherules appears to overlap with eGFP (Figure 5D). Conversely, in SINV 550C-infected cells, spherule eGFP intensity values displayed a broader unimodal distribution, indicating much more eGFP leakage into spherules (Figure 5D). Therefore, we conclude that, in SINV 550C-infected cells, few viral replication spherules exclude eGFP, whereas most do not. We also found a positive correlation between J2 intensity and eGFP intensity within individual spherules in wild-type SINV-infected *Dcr2*+ cells (Figure 5E). This correlation was even stronger in SINV 550C-infected cells, suggesting that impaired spherule integrity is linked to higher levels of processed dsRNA in a subset of spherules (Figure 5F).

### The SINV Opal-to-Cys variant induces a higher interferon response in human cells

Our previous study showed that the nsP-processing defects of the SINV 550C variant also occur in vertebrate cells (*1*). Unlike in insects, RNAi is not the primary immune defense pathway in vertebrate cells. Instead, host-encoded pattern recognition receptors (PRRs), RIG-I and MDA5, detect viral dsRNA and trigger downstream interferon (IFN) responses (*39*, *40*). Based on our findings that nsP-processing delays cause spherule integrity defects in mosquito cells, we reasoned that similar spherule defects occur during SINV 550C infection in vertebrate cells, thereby activating dsRNA sensors and inducing innate immune responses. If so, we would expect SINV 550C-infected cells to trigger a stronger immune response than wild-type SINV (550opal)-infected cells.

We tested this hypothesis by infecting human A549 cells with either SINV 550C or SINV 550opal for 16 hours, then monitoring changes in immune gene expression using bulk RNA sequencing. Consistent with our expectations, we observed increased expression of several antiviral interferon-stimulated genes (ISGs) and genes related to Type I/III IFN responses in SINV 550C-infected cells (Figure 6A-B), indicating that the loss of the opal codon may result in higher IFN production and signaling in vertebrate cells.

**Figure 6.**
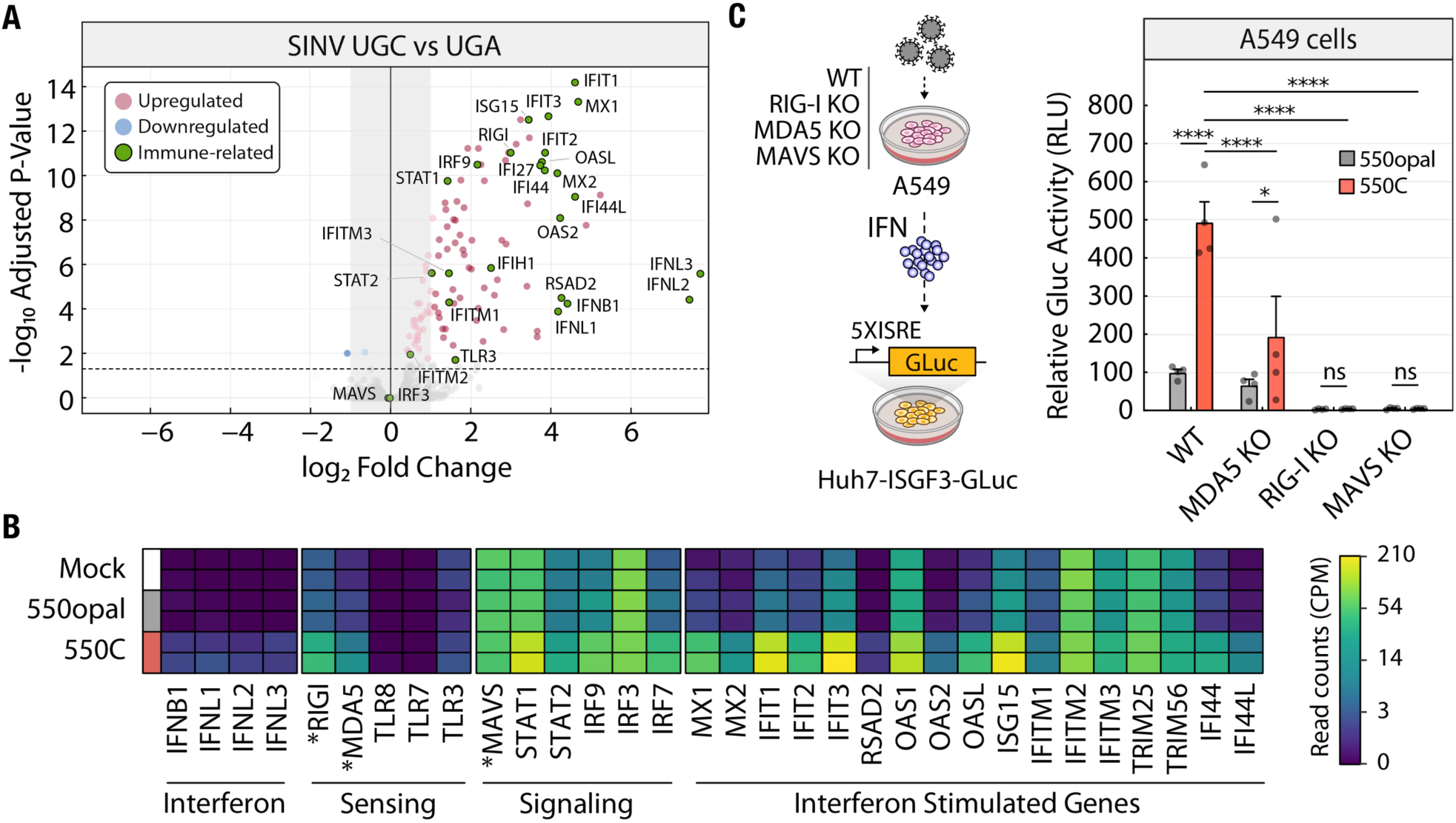
The SINV sense-codon variant induces a higher immune response in human cells. **(A)** Volcano plot of differentially expressed genes between A549 cells infected with wild-type (550opal) and variant (550C) SINV. ISGs and other immune-related genes are represented as green circles. **(B)** Changes in immune-associated gene expression 16 hours post-infection with wild-type (550opal) and variant (550C) SINV in human A549 cells. RLR and MAVS expression are highlighted with asterisks (*). **(C)** Experimental workflow to measure IFN production in A549 cells infected with either wild-type SINV 550opal or SINV 550C at an MOI of 5. Huh7 cells expressing a 5XISGF3-GLuc reporter were treated for 24 h with A549 cell supernatants, and secreted Gaussia luciferase was quantified. Reporter activity in 5XISGF3-GLuc Huh7 cells treated with supernatants collected 16h post-infection from wild-type, *MDA5* KO, *RIG-I KO*, or *MAVS* KO A549 cells. The data represent the mean of four independent biological replicates. Two-way ANOVA with Sidak’s multiple comparisons test. **** = p < 0.0001, * = p < 0.05, ns = not significant.

To test this, we measured IFN production from infected A549 cells in Huh7 cells expressing a 5XISGF3-GLuc reporter (Figure 6C left) (*41*). Consistent with the transcriptomic response, we observed significantly higher GLuc activity in cells treated with supernatants from SINV 550C-infected A549 cells than in those treated with supernatants from wild-type SINV (550opal)-infected A549 cells (Figure 6C right).

A549 cells respond to dsRNA by upregulating RLRs (Retinoic acid-inducible gene I (RIG-I)-like receptors) – RIG-I and MDA5 (*42*). Both RLRs signal through the downstream adapter MAVS (Mitochondrial Antiviral Signaling protein) and are crucial for IFN induction during alphavirus infection (*40*). After confirming elevated expression of RIG-I and MDA5 in SINV 550C-infected cells (Figure 6B), we asked whether these RLRs are functionally important for immune stimulation in response to SINV 550C infection by quantifying IFN production in their presence and absence (Figure 6C right). First, we asked whether MDA5, the cytoplasmic RNA sensor that detects long double-stranded RNA (dsRNA), is responsible for elevated interferon production. We infected MDA5 KO A549 cells with either SINV 550C or wild-type SINV (550opal) and measured the interferon response at 16 hours post-infection (hpi). We found that MDA5 loss significantly reduced, but did not ablate, IFN production in SINV 550C-infected A549s (Figure 6C right). However, loss of RIG-I (RIG-I KO) or the adapter MAVS (MAVS KO) led to near-complete loss of IFN production in SINV 550C- and wild-type SINV (550opal)-infected A549s (Figure 6C right). These results suggest that human RLRs, primarily RIG-I and MDA5, contribute to ligand detection when viral spherule integrity is compromised in SINV-infected human cells.

We next asked whether the S.A. AR86-specific nsP1 I538T mutation alters the SINV 550C-induced immune response. The Thr (T) residue at nsP1 position 538 has previously been shown to be necessary and sufficient to prevent immune activation in S.A. AR86 by inhibiting JAK/STAT signaling in murine fibroblasts (*43*, *44*). Indeed, cells infected with SINV 550C carrying the I538T mutation produced less IFN than SINV 550C (Figure S13). However, consistent with our earlier observation in RNAi-competent mosquito cells, which suggests that an I538T-induced delay in nsP processing may also promote IFN induction, we observed higher IFN levels in cells infected with SINV carrying an I538T mutation relative to wild-type SINV (550opal) (Figure S13).

Overall, our data support a model in which delayed nsP processing, caused by an Opal-to-Cys substitution at the stop codon between nsP3 and nsP4, produces defective or incomplete spherules in alphavirus-infected cells (Figure 7). These structurally compromised spherules allow cytoplasmic proteins, including RNA sensors like Dcr2 in insect cells and RLRs RIG-I and MDA5 in vertebrate cells, to access viral dsRNA. This access enhances immune activation and ultimately lowers viral fitness in immune-competent insect and vertebrate cells (Figure 7B). Therefore, the nearly universally conserved nsP3 opal codon in alphaviruses is crucial for hiding viral RNA replication and preventing immune activation in both insect and vertebrate hosts.

**Figure 7.**
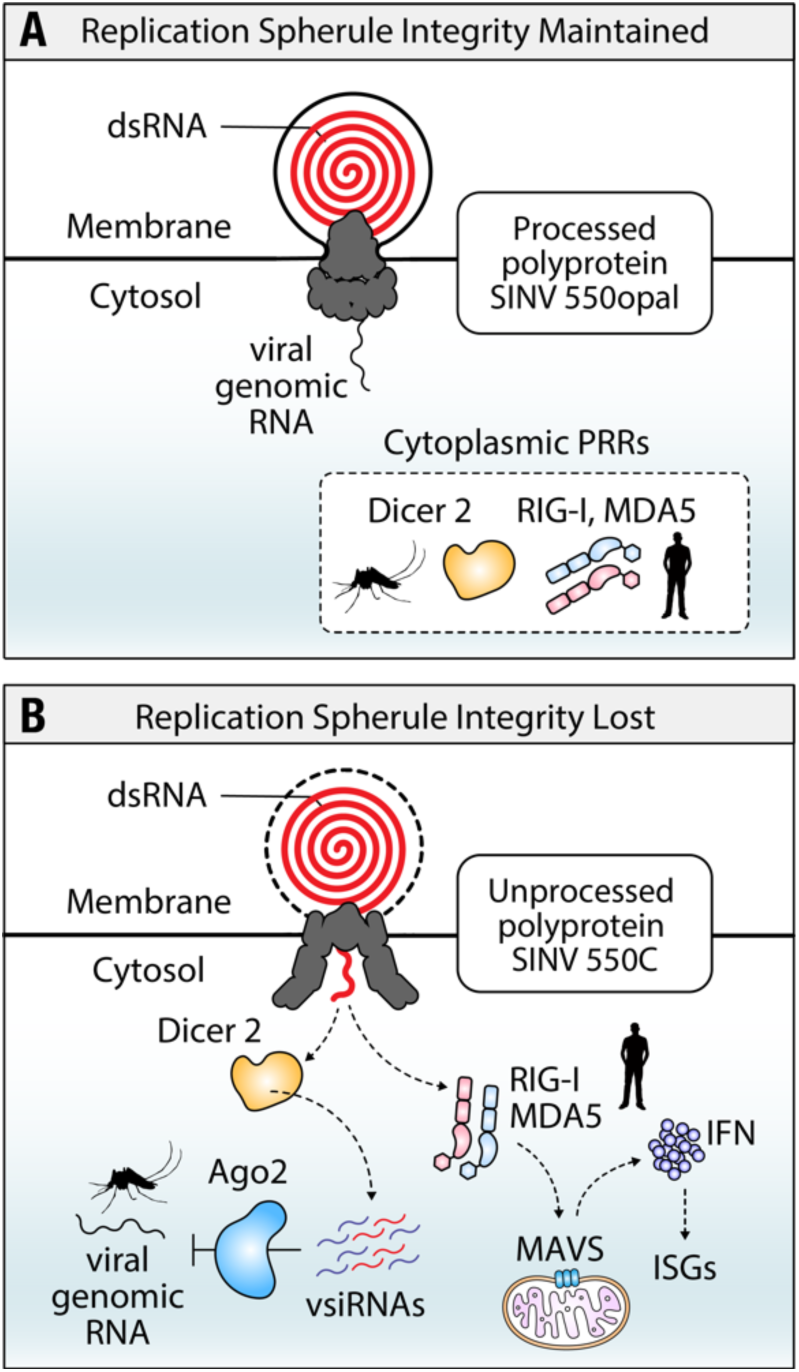
Proposed model of how opal-to-sense codon substitutions reduce viral fitness. **(A)** The SINV nsP3 opal codon helps maintain replication spherule integrity to avoid detection of viral RNA by cytoplasmic RNA nucleases (Dcr2) or sensors (RIG-I, MDA5) in mosquito or vertebrate cells, respectively. **(B)** Improper processing of the nsP polyprotein in SINV 550C variant-infected cells leads to the loss of spherule integrity, resulting in excessive Dcr2- and Ago2-mediated antiviral RNAi induction in mosquito cells and RIG-I /MDA5-mediated, MAVS-dependent IFN induction and ISG expression in vertebrate cells.

## Discussion

Our study demonstrates that retaining the highly conserved nsP3 opal stop codon is a multipotent defense strategy used by alphaviruses (Figure 7A). By systematically dissecting its function in both insect and vertebrate cells, we find that the opal codon is crucial not only for a temperature-dependent trade-off between nsP production and processing, as we previously showed (*1*), but also for maintaining the physical integrity of replication spherules, thereby shielding viral dsRNA from host immune sensors. We find that opal-to-sense substitutions (e.g., SINV 550C) disrupt the cadence of alphavirus non-structural polyprotein processing, leading to spherules with compromised integrity and increased exposure of viral dsRNA (Figure 7B). This defect enables cytoplasmic Dcr2 in mosquitoes to access and process viral dsRNA, triggering a robust antiviral siRNA response that sharply reduces viral fitness in RNAi-competent *Aedes albopictus* cell lines and *Aedes aegypti* mosquitoes *in vivo* (Figures 1B, 2C-G, 7B). Using small RNA sequencing, we show that SINV 550C, which differs from wild-type SINV by a single nucleotide at the opal codon, induces a significantly higher small RNA response in *Aedes albopictus* cells (Figure 3B). This direct measurement of RNAi activity in response to wild-type and variant SINV infections provides conclusive evidence for the role of RNAi in the fitness advantage conferred by the opal codon. These findings extend earlier work in *Anopheles gambiae*, where an Opal-to-Arginine mutation reduced *in vivo* ONNV infectivity, underscoring the evolutionary importance of preserving the opal codon for efficient vector transmission (*4*). We also show that alphaviral structural RNA elements, such as E1-hs, protect viral fitness against host RNAi by acting as ‘decoy’ hotspots of vsiRNA production, likely blunting the antiviral RNAi response (Figure 4C-G). Thus, alphaviruses use a two-pronged defense strategy against mosquito RNAi: first, by physically sequestering dsRNA within replication spherules, and second, by producing ‘decoy’ vsiRNAs from structured genomic RNA elements. Beyond insects, compromised spherule integrity in human cells may also permit RNA sensors, such as RIG-I and MDA5, to detect viral dsRNA, thereby enhancing interferon signaling and ISG expression (Figures 6 and 7B).

Recent *in situ* structural and functional characterization has provided insights into the architecture of mature alphavirus replication spherules – membrane-associated structures that shield viral dsRNA from cytoplasmic RNA sensors in vertebrate and mosquito cells (*28*, *45–48*). Although mature spherule biogenesis remains poorly understood, it is thought to require proteolytic processing of P1234 into nsP1-4 proteins that comprise the ‘neck’ of the replication spherule (*2*, *35*, *45*, *49*, *50*). Using quantitative immunoblotting and confocal microscopy, we show that incomplete polyprotein processing in SINV 550C disrupts the structural integrity and permeability of replication spherules, potentially allowing cytoplasmic Dcr2 access to viral dsRNA in infected mosquito cells (Figures 5, S11). Additionally, the J2 signal intensity in SINV 550C-infected cells exceeds that in wild-type SINV (550opal)-infected cells, but only in the presence of Dcr2, suggesting that the excess J2 signal may represent Dcr2-processed viral dsRNA and thereby contribute to the overall increase in the siRNA response (Figures 5C, S12B-C). Therefore, increased vsiRNA induction by SINV 550C in *Dcr2*+ cells correlates with slower processing of viral nsP (Figure S11). Although it is clear that the opal-to-Cys mutation increases the permeability of viral spherules, the directionality of dsRNA sensing remains unclear. Future studies, aided by new tools and advances in cryo-electron tomography, could investigate whether the higher RNAi induction in SINV 550C-infected mosquito cells is due to Dcr2 gaining access to the viral replication compartment or to viral dsRNA leaking from spherules into the host cytoplasm (*45*, *51*).

Together, these findings suggest that although some opal-to-sense substitutions may promote viral replication in Dcr2-deficient mosquito cells, the resulting delay in polyprotein processing may compromise the integrity of replication spherules. This structural vulnerability increases viral susceptibility in the presence of Dcr2, ultimately limiting the replication efficiency of sense-codon variants in Dcr2-competent cells. Our results align with previous findings of a Dcr2-dependent change in the relative replication of CHIKV replicons carrying mutations that delay nsP processing, potentially indicating a generalizable trade-off between replication efficiency and RNAi susceptibility (*52*).

Because RNAi is the primary antiviral defense in mosquitoes, mosquito-transmitted viruses have evolved mechanisms to circumvent or suppress the RNAi response while maintaining persistent, non-pathogenic infections in their vectors. For example, non-coding subgenomic flaviviral RNAs (sfRNAs) at the 3’ end of flavivirus genomes outcompete native Dcr2 and Ago2 substrates for binding, thereby reducing RNAi activity (*53*). Other viruses, such as Flock House virus (FHV B2, *Nodaviridae)* and Culex Y virus (CYV VP3, *Birnaviridae),* encode *bona fide* viral suppressors of RNAi (VSRs) (*54–56*). Until recently, alphaviruses were not known to encode sfRNAs or proteins with strong VSR activity. In fact, SINV expressing heterologous FHV B2 exhibits high vector mortality and reduced virus transmission (*57*). Thus, unlike other arboviruses, alphaviruses likely avoid strong, direct suppression by RNAi, instead dampening or evading the mosquito RNAi response to maintain persistent, non-lethal infection in insect hosts. However, some alphavirus proteins exhibit VSR activity, most notably CHIKV nsP2/3 and SFV capsid, which weakly suppress RNAi in mammalian cells, potentially by sequestering viral dsRNA (*58*, *59*). More recent work provides compelling evidence that mature SINV nsP2 also antagonizes mosquito RNAi using a similar mechanism (*60*). Notably, cleavage-deficient SINV nsP2 lacks VSR activity, lowering *in vivo* fitness in immune-competent mosquitoes. Thus, it is formally possible that lower levels of processed nsP2, caused by slower polyprotein processing, also contribute to lower SINV 550C fitness.

One intriguing model for how alphaviruses suppress mosquito RNAi comes from a study of Semliki Forest Virus (SFV), which is unusual among alphaviruses in that it predominantly lacks the nsP3/4 opal stop codon (*32*). SFV genomes produce ‘decoy’ vsiRNAs during infection of mosquito cells that swamp the RISC machinery, thereby protecting RNAi-susceptible viral targets. These vsiRNAs arise from genomic hotspots that correspond to structured RNA regions but are not antiviral themselves. Consistent with this ‘decoy’ vsiRNA model, we identified a vsiRNA hotspot in SINV that maps to a structured RNA element within the E1 coding region (Figures 4A-C). Increased RNA exposure in SINV 550C-infected cells disproportionately reduces vsiRNA production from this hotspot element (Figure 4A-inset). Disrupting this RNA secondary structure (E1-hs) significantly reduced SINV 550C replication in RNAi-competent cells, underscoring its role in RNAi evasion (Figures 4D-G). Structural conservation of this RNA element also suggests that it may perform a similar function in other alphaviruses (Figures S9 and S10). Our data show that such decoy strategies can be generally beneficial but are especially critical when viruses are more susceptible to Dcr2 inhibition due to defects in spherule integrity, which might be caused by genetic mutations (as we show in this study) or by other biophysical changes within host cells (Figures 4).

Our results also demonstrate a link between alphavirus nsP processing and innate immune detection in vertebrate cells. Impaired replication spherule integrity in SINV 550C increases IFN production and antiviral ISG expression by enhancing recognition by the cytoplasmic RNA sensors RIG-I and MDA5 in human A549 cells (Figure 6) (*40*). Furthermore, a mutation (nsP1 I538T) associated with slower P1/2 processing and a ‘double processing delay,’ in combination with an Opal-to-Cys substitution, also leads to higher IFN levels (Figure S13). Our findings align with recent studies showing that alphaviruses with delayed nsP processing mutations elicit stronger immune responses by generating host-derived rPAMP RNAs that serve as RLR ligands (*39*, *61*, *62*). Therefore, host-derived dsRNAs might also promote immune activation in SINV 550C-infected cells. Although our data cannot distinguish between viral- and host-derived PAMP RNAs as the source of stimulation, it is clear that IFN-induction relies on detection by the RLRs RIG-I and MDA5 (Figure 6C).

The 538Thr residue in SINV nsP1 has been shown to suppress IFN production and antagonize downstream IFN signaling, thereby increasing neurovirulence in mice (*63*, *64*). Surprisingly, however, we did not observe strong IFN suppression by SINV nsP1 538Thr in human cells. Although we cannot rule out the possibility that the SINV nsP1 I538T mutation might have preexisted in the original mosquito-isolated S.A. AR86 strain, it is just as likely that this mutation arose during extensive passaging (> 45 passages) in mice (*65*). Therefore, based on our current data, we conclude that its immunosuppressive phenotype may be specific to mouse cells (*43*, *65*).

Collectively, the results presented in this study extend our previous model, in which the alphavirus nsP3 opal codon helps balance polymerase (RdRp) production and nsP processing efficiency (*1*). We now demonstrate that the opal codon also serves an immune-specific function by enabling alphaviruses to evade triggering an antiviral response (Figure 7). This may also explain why the opal codon is strictly retained across insect-restricted alphaviruses that do not face vertebrate- or temperature-specific constraints described in our previous study (*1*).

One of the central conflicts faced by viruses infecting evolutionarily divergent host organisms is that adaptations that confer fitness benefits in one host, such as vertebrates, may be deleterious in another, such as mosquitoes. Some viral adaptations might confer benefits in both hosts. Our study highlights one adaptation – the nsP3 opal codon – that confers benefits in both hosts by maintaining replication spherule integrity and shielding viral RNA from cytoplasmic nucleases and RNA sensors (Figure 7). In doing so, this single codon confers a fitness advantage in RNAi-competent mosquito cells while also limiting interferon responses in mammalian cells. Thus, the retention of the opal codon by alphaviruses reflects evolutionary pressure from immunity in both hosts. By demonstrating that the alphavirus opal codon is a multipurpose defensive adaptation, our work reframes a classical paradigm of alphavirus biology and opens new avenues for research into vector-borne disease control.

## Materials and Methods

### Insect and Mammalian Cell Culture

C6/36 *Aedes albopictus* cells were grown at 28°C under 5% CO_2_ in humidified incubators and cultured in high-glucose, L-glutamine Minimal Essential Medium (Gibco) supplemented with 10% fetal bovine serum (Cytiva) and 1% penicillin-streptomycin (Gibco). U4.4 *Aedes albopictus* cells were grown at 28°C under 5% CO_2_ in humidified incubators. They were cultured in high-glucose, L-glutamine Mitsuhashi and Maramorosch Insect Medium w/o Sodium bicarbonate (VWR) supplemented with 0.12 gm/L Sodium bicarbonate, 10% fetal bovine serum (Cytiva), and 1% penicillin-streptomycin (Gibco). The *Aedes albopictus* U4.4 cells were a gift from Dr. Douglas (Doug) Brackney (University of Connecticut).

### CRISPR editing of Dicer 2 in U4.4 cells

5X10^5^ U4.4 cells were seeded into 6-well plates and co-transfected with 6.25 µg of purified SpyCas9-NLS (PNA Bio #CP01-50) and a synthesized single guide RNA (sgRNA) (AUAUUCGACGAAUGUCACCA) targeting the 5’ end of the *Dcr2* gene (Synthego). The guide RNA was designed using CHOP-CHOP and verified with Cas-OFFinder to assess off-target effects by searching the guide RNA sequence against the *Aedes albopictus* genome (JXUM01). The synthesized sgRNA was also modified with 5’ 2’-O-methyl (OMe) analogs and 3’ phosphorothioate (PS) to enhance stability and performance. We harvested 1X10^6^ cells two weeks after Cas9-sgRNA transfection, extracted gDNA, and amplified the sgRNA target region using flanking primers (see Data S2 for information regarding primers used in the study). Purified PCR products were sent for Sanger sequencing, and the resulting chromatograms were used for TIDE analyses (version 5.0), which revealed a knockout efficiency of 82-85% (Figure S2) (*66*).

### Ago2 silencing in U4.4 cells

We used dsRNA to silence *Aedes albopictus* Ago2 in U4.4 cells, as previously described (*38*). Briefly, total RNA from U4.4 cells was reverse transcribed with oligo(dT) to synthesize cDNA. The cDNA served as the template for PCR to amplify a region within the PIWI domain of Ago2 with primers carrying T7 promoters at the 5’ end (see Data S2 for information regarding primers used in the study). Purified PCR products were used as templates for in vitro transcription with T7 RNA polymerase (NEB) to generate double-stranded RNA (dsRNA), which was then purified using the MEGAclear clean-up kit (Invitrogen) according to the manufacturer’s instructions. 5X10^5^ U4.4 cells were transfected with 500ng of Ago2 dsRNA using Lipofectamine 3000 (Invitrogen), and Ago2 silencing was assessed over 1-4 days by qRT-PCR (see Data S2 for information regarding primers used in the study). To determine the extent of functional disruption of the siRNA pathway following Ago2 silencing, we used a previously described eGFP-reporter silencing assay with dsRNA targeting eGFP (dsGFP) (Figure S5A) (*58*). Infections were performed 48 hours after Ago2 dsRNA treatment.

### Small RNA sequencing and analysis

10 µg of total RNA was isolated from cultured U4.4 and C6/36 mosquito cells five days after SINV infection. RNA purity and integrity were assessed using the Agilent 4200 TapeStation. All subsequent small RNA enrichment steps were performed by Novogene, USA, using 10 µg of total RNA. Small RNA sequencing was performed on an Illumina NovaSeq (Novogene, USA). Three biological replicate libraries were sequenced for each sample. After removing adaptor sequences (Trimmomatic), the processed reads were mapped to the SINV genome (strain TE12) using Geneious Mapper with a mapping quality of 30, a 10% gap, and a 20% mismatch rate. The standard counts-per-million (CPM) method (CPM = gene read count/total mapped reads × 1,000,000) was used to normalize for sequencing depth when comparing small RNA counts between samples. Logoplots showing nucleotide frequencies within vpiRNA reads were generated from 24-31nt vpiRNA reads and visualized using WebLogo 3 (*67*).

### Virus Constructs

All viral stop-codon mutant plasmids were generated by site-directed mutagenesis. SINV sense-codon variants were generated in our previous study (*1*). The SINV E1-hs_mut_ variant was generated using the following primer pair (see Data S2 for information regarding primers used in the study). Mutagenic primers were designed with Takara’s In-Fusion Cloning Primer Design tool. PCR reactions were performed with Phusion polymerase (NEB) using the following cycling conditions: 1. 98°C for 30 sec, 2. 98°C for 30 sec, 3. 50°C for 30 sec, 4. 72°C for 7 min (repeat steps 2-4 18x), 5. 72°C for 10 min, 6. Hold at 4°C. Reactions were then treated with *DpnI* for 6 hours and purified with the Monarch PCR and DNA Clean-up Kit (NEB) according to the manufacturer’s protocol. The purified reaction was transformed into DH5α cells, and DNA was isolated with the Monarch Plasmid Miniprep Kit (NEB) according to the manufacturer’s protocol. Luciferase-based reporter viruses were generated by digesting parental and mutant plasmids with *SpeI* (NEB) and cloning in luciferase (Nano) using HiFi Gibson assembly (NEB). The Gibson reactions were then transformed into DH5α cells, and DNA was isolated with the Monarch Plasmid Miniprep Kit (NEB) according to the manufacturer’s protocol.

### Independent Viral Growth Assays

2 µg of infectious clones were linearized with *XhoI* (NEB) and subjected to *in vitro* transcription (IVT) with SP6 RNA polymerase (NEB) according to the manufacturer’s protocol. Cells were seeded into 24-well plates to reach 70-80% confluence and transfected with IVT using Lipofectamine LTX (Thermo Fisher) according to the manufacturer’s protocol. Cells were collected 48 hours post-transfection and analyzed by flow cytometry (BD Fortessa). Infection rates for all variants were normalized to wild-type SINV (550opal).

### Viral Competition Assays

Genome copy numbers of WT and mutant virus stocks were quantified by RT-qPCR (see Data S2 for information regarding primers used in the study). Briefly, cDNA synthesis was performed on viral supernatant using M-MuLV Reverse Transcriptase (NEB) with oligo(dT) (20mer + 5’-Phos) (IDT) according to the manufacturer’s protocol. RT-qPCR was performed using the SYBR Green Master Mix (Thermo Fisher) with gene-specific primers on the Applied Biosystems StepOnePlus Real-Time PCR System (Life Technologies) according to the manufacturer’s protocol. WT and mutant SINV virus stocks were then normalized to equal genome copy numbers. Wild-type and *Dcr2* KO U4.4 cells, and C6/36 cells were seeded into 24-well plates and infected with TE3’2J-mCherry WT virus alongside TE3’2J: eGFP WT or mutant viruses at a 1:1 ratio (MOI – 0.1). Cells were collected 48 hours post-infection and analyzed by flow cytometry (BD Fortessa) to determine the percentage of cells infected with WT (red) and variant (green) viruses. The viral competitive index was calculated as the ratio of variant (eGFP)-to-WT (mCherry)- infected cells within the population.

### Generation of trans-heterozygous Dcr2 mutant mosquitoes

#### Dcr2 eGFP null line

We previously developed a transgenic line containing eGFP, as originally described in Basu et al. (*21*). Initially, we created an out-of-frame deletion using TALEN technology. Loss of function was confirmed by crossing into our “sensor” line and observing disease phenotypes after challenge with alphavirus and flavivirus pathogens (*22*, *68*). Using the same TALEN target site, we knocked in a construct containing eGFP under the Poly-UB promoter, as described in Basu et al. (*21*).

#### Dcr2 dsRED null line

For insertion of the dsRED-containing construct, we employed sgRNA-directed Cas9 nuclease. The sgRNA was designed to target a site adjacent to the original TALEN site. We developed two distinct donor constructs with different homology arm sequences. Before creating these constructs, we sequenced the target site, revealing polymorphisms in the intronic regions of the homology arms. To account for this genetic variation, we generated donor constructs based on the two most prevalent homology arm sequences identified across individual mosquito specimens. Both donor constructs were co-injected during the transformation procedure. We confirmed one insertion site junction, but the second remains unverified. However, confirming genomic insertions in *Aedes aegypti* presents significant technical challenges due to the genome’s architecture and heterogeneity, particularly the presence of numerous large introns containing many repetitive sequences and the previously mentioned polymorphisms.

*Dcr2* homozygous knockouts: We generated trans-heterozygous *Dcr2*−/− mosquitoes by crossing two distinct Dcr2 mutant lines, one expressing eGFP (whole-body) and the other expressing dsRED (whole-body) (Figure 2A). The progeny from these crosses were screened at the larval stage for fluorescent marker expression.

### In vivo mosquito infections

Adult *Aedes aegypti* mosquitoes were infected by injecting 0.5 µl of Sindbis virus (SINV; 10^6^ TCID_50_/ml) diluted in Dulbecco’s Modified Eagle Medium (DMEM) directly into the thorax. Wild-type siblings, identified as negative for eGFP and dsRED fluorescence, served as controls. After inoculation, *Dcr2* null (*Dcr2* −/−) mutants and wild-type siblings were maintained at 28 °C and 80% relative humidity under a 14:10 h light: dark cycle. Mosquitoes were collected at 0 and 72 hours post-infection in pools of five, flash-frozen in liquid nitrogen, and stored at −80 °C until further analysis. Quantitative RT-PCR (qRT-PCR) was performed to determine viral RNA levels in total RNA extracted from pools of five mosquitoes per replicate. A separate set of individual mosquitoes (n = 15/condition) was collected and flash-frozen for quantification of infectious virus titer via TCID_50_ assays on Vero cells. Mosquitoes were homogenized in 60 µL of sterile 1× PBS containing protease inhibitor, and the homogenate was clarified by centrifugation through a 0.45 µm Ultra-free PVDF filter. Survival curves for wild-type SINV (550opal) we*r*e generated over 22 days for ≥28 female mosquitoes infected with 10^6^ TCID_50_/ml wild-type SINV, comparing *Dcr2*+/+ and *Dcr2*−/− mosquitoes.

### Total RNA Extractions and Real-Time Quantitative RT-PCR Analysis

To quantify minus- and plus-strand RNA copies, cells were seeded into a 24-well plate and infected with WT or variant strains at greater than 90% confluence. At 4 and 18 hours post-infection, cells were harvested with TRIzol reagent (Thermo Fisher), and RNA was extracted with the Direct-zol RNA Miniprep kit (Zymo) according to the manufacturer’s protocol. Following RNA extraction, cDNA was synthesized with M-MuLV Reverse Transcriptase (NEB) using strand-specific primers or OligoDT (20mer+5’Phos) (IDT) according to the manufacturer’s protocol. In all cases, RT-qPCR was performed with SYBR Green master mix (Thermo Fisher) and plus-strand-specific primers on the Applied Biosystems StepOnePlus qRT-PCR machine (Life Technologies) (see Data S2 for primer details).

### Bulk RNA sequencing

To quantify global transcriptomic changes, human A549 cells were infected with wild-type (550opal) and variant (550C) SINV at an MOI of 5. Mock- and virus-infected cells were collected 16 hours post-infection in Zymo DNA/RNA buffer and sent to Plasmidsaurus for bulk RNA sequencing. Analysis was carried out by Plasmidsaurus using the following method: Quality of the fastq files was assessed using FastQC v0.12.1. Reads were then quality-filtered using fastp v0.24.0 with poly-X tail trimming, 3’ quality-based tail trimming, a minimum Phred quality score of 15, and a minimum length of 50 bp. Quality-filtered reads were aligned to the reference genome using STAR v2.7.11 with non-canonical splice junction removal and output of unmapped reads, followed by coordinate sorting with Samtools v1.22.1. PCR and optical duplicates were removed using UMI-based deduplication with UMIcollapse v1.1.0. Alignment quality metrics, strand specificity, and read distribution across genomic features were assessed using RSeQC v5.0.4 and Qualimap v2.3, and results were aggregated into a comprehensive quality control report using MultiQC v1.32. Gene-level expression quantification was performed using featureCounts (subread package v2.1.1) with strand-specific counting, multi-mapping read fractional assignment, exons, and three prime UTR as the feature identifiers, and grouping by gene_id. Final gene counts were annotated with gene biotype and other metadata extracted from the reference GTF file. Differential expression was performed with edgeR v4.0.16 using standard practice, including filtering for low-expressed genes with edgeR::filterByExpr using default values.

### Viral Replication Assays

SINV nsP3-Nluc reporter viruses were generated by digesting 2 µg of plasmid DNA with *XhoI* and performing in vitro transcription (IVT) with SP6 RNA polymerase (NEB) according to the manufacturer’s protocol. Cells were seeded into black-walled, clear-bottom 96-well plates at 70-80% confluency and transfected with 100 ng of capped viral RNA using Lipofectamine LTX (Thermo-Fisher) according to the manufacturer’s protocol. At 48 hours post-transfection, translation was quantified using the NanoGlo luciferase assay system (Promega) according to the manufacturer’s protocol. Luminescence was measured using a Cytation3 Imaging Reader (BioTek).

### IFN reporter activity

To quantify secreted IFN levels in infected wild-type, *MDA5* KO, and *MAVS* KO A549 cells, cell supernatants from wild-type SINV (550opal) and 550C-infected cells were collected 16 hours post-infection and transferred to 5XISGF3-GLuc Huh7 reporter cells that stimulate production of Gaussia luciferase downstream of a 5XISRE reporter that is induced by ISGF3 (*41*). Reporter cells were incubated at 37°C and 5% CO_2_ for 48 hours to allow GLuc secretion into the media. Secreted GLuc was quantified using the Pierce Gaussia Luciferase Assay kit (Thermo Fisher Scientific) on a Cytation3 plate reader (BioTek) according to the manufacturer’s protocol. The 5XISGF3-GLuc Huh7 reporter cells were a gift from Dr. Ram Savan (University of Washington). The *RIG-I KO, MDA5 KO,* and *MAVS* KO A549 cells were a gift from Dr. Dan Stetson (University of Washington).

### Western Blot Analyses

Cells were seeded into six-well plates and infected at an MOI of 5. After a 1-hour incubation at 4°C, cells were washed with cold 1X PBS, and lysates were harvested in RIPA buffer (Pierce) supplemented with 1X protease inhibitor (cOmplete). Purified protein was denatured in 2X Laemmli buffer (BioRad) with 5% β-Mercaptoethanol (Sigma-Aldrich). Immunoblots were run on Mini-TGX precast gels and transferred to a Trans-Blot 0.2 µm nitrocellulose membrane using the Trans-Blot Turbo Transfer System (Bio-Rad). Blots were incubated overnight at 4 °C in 5% BSA with the primary antibody at a 1:3,000 dilution, then washed with 1X TBS containing 0.1% Tween-20. Blots were probed with a secondary α-rabbit HRP-conjugate (R&D Systems) and visualized using a Bio-Rad ChemiDoc Imaging System. Immunoblots probing for non-structural protein expression and polyprotein processing were performed with independent biological replicates for each virus under different host conditions.

### Immunofluorescence Confocal Microscopy

Glass-bottom #1.5 coverslip 12-well plates (NC0190134; Thermo Fisher Scientific) were treated with 200 µl of sterile 0.01% poly-L-lysine (P4832; Sigma-Aldrich) per well for 1 hour at room temperature, rinsed once with DPBS++ (14-040-216; Thermo Fisher Scientific), and allowed to dry for 1 hour. U4.4 cells were seeded onto poly-L-lysine-coated wells and grown to approximately 20% confluency. Cells were then infected with wild-type (WT) or 550C SINV 3XF-GFP at a high multiplicity of infection (MOI) and incubated for 48 hours. Cells were rinsed twice with DPBS++ and fixed with 200 µl of 4% (v/v) paraformaldehyde (28906; Thermo Fisher Scientific) in DPBS++ for 10 minutes, followed by two washes. Permeabilization was performed with 400 µL of 0.05% Triton X-100 (BP151; Thermo Fisher Scientific) in DPBS++ for 5 minutes, followed by two washes. Cells were then blocked with 400 µl of 10% (w/v) bovine serum albumin (BSA; BP9703100; Thermo Fisher Scientific) in DPBS++ for 1 hour.

Steps were performed in the dark to minimize photobleaching. Primary antibody staining was performed with 500 µl of mouse monoclonal anti-dsRNA J2 (1:1000) (76651L; Cell Signaling) in 5% BSA, incubated overnight at 4°C with gentle agitation. Cells were washed three times with DPBS++ and blocked again with 400 µl of 10% BSA in DPBS++ for 1 hour. Secondary antibody staining was performed with 500 µl of Alexa Fluor 633-conjugated goat anti-mouse antibody (1:1000) (A-21050; Invitrogen) in 5% BSA, and the sample was incubated overnight at 4°C without agitation. After a single wash with DPBS++, the nuclei were counterstained with Hoechst 33258 (1:1000) (H1398; Invitrogen) in 5% BSA for 15 minutes, then washed three times. Coverslips were mounted with 150 µl of SlowFade™ Diamond Antifade Mountant (S36972; Thermo Fisher Scientific). All steps were performed at room temperature unless otherwise noted. Each wash was performed with DPBS++ for 5 minutes. Imaging was performed on a Leica STELLARIS confocal microscope (DMi8 stand) with a 63x oil-immersion objective and Leica Application Suite X (LAS X) software. All comparative microscopy images were acquired on the same day with identical settings.

### Microscopy Imaging Analysis

Images were analyzed using FIJI/ImageJ 1.54f with Java 1.8.0_322 (64-bit) and the ImageJ 3D Suite MCIB V3.96 (*69–71*). Spherule boundaries were determined by MaxEntropy thresholding of the anti-dsRNA Alexa Fluor 633 signal and used to create 3D regions of interest (ROIs). The accessibility of spherules to cytoplasmic eGFP was quantified as the mean eGFP signal intensity within entire spherules using the 3D Manager Quantif3D function in the ImageJ 3D Suite plugin. Spherule boundaries (Figure S12D) were determined by MaxEntropy thresholding of the anti-dsRNA Alexa Fluor 633 signal and used to create 3D regions of interest (ROIs). Reported mean eGFP signal intensities are per-spherule. Cellular dsRNA levels were quantified as the mean anti-dsRNA Alexa Fluor 633 signal using the 3D Manager Quantif3D function in the ImageJ 3D Suite plugin. Infected cells were identified by the presence of cytoplasmic eGFP and dsRNA. Cellular boundaries were determined by Otsu thresholding of the cytoplasmic eGFP signal, and areas excluding eGFP were filled using the binary ‘Fill in Holes’ function, creating 3D ROIs as defined by the 3D Object Counter with a minimum threshold of 1,500 cubic voxels. If needed, neighboring cells were separated across the entire stack before quantification using the 3D Watershed function in the ImageJ 3D Suite plugin. Nuclei were used as seeds. Reported mean J2 signal intensities are per cell.

### Phylogenetics and RNA Structure Analysis

A multiple sequence alignment of nsP4 coding sequences from 49 extant alphavirus species was generated with Clustal Omega. The alignment was manually curated in Geneious, and a maximum-likelihood phylogenetic tree was inferred in PHYML using the HKY85 substitution model with 100 bootstrap replicates for statistical support. The tree was visualized with FigTree. RNA secondary structure conservation was assessed using RNAalifold on a multiple-sequence alignment of the 45-nt E1-hs region from 2,086 alphavirus sequences collected from BV-BRC (v3.51.7) (*72*). SHAPE-constrained RNA secondary structure was generated with Vienna RNAfold using SHAPE-MaP data previously reported by *Kutchko et al.* and Madden et al. (*30*, *31*).

### Statistics

Statistical analyses were performed using GraphPad Prism (v10.2, GraphPad Software Inc., San Diego, CA). Data were first assessed for normality (Gaussian distribution) using the Kolmogorov-Smirnov test. For data meeting normality (α = 0.05) and homogeneity of variance criteria, means were compared using a Two-Way ANOVA (for multiple groups with two variables), unless otherwise stated, with post hoc tests for multiple comparisons. Some data were log-transformed prior to statistical testing to satisfy assumptions of normality. In these cases, tests were performed on the transformed values.

## Acknowledgments

We thank the members of the Malik and Emerman labs for their valuable discussions and acknowledge Megan Levy, Peter Dietzen, and Rebecca Fereira Alves for their feedback on the manuscript. We thank Richard W. Hardy and Suchetana (Tuli) Mukhopadhyay (Indiana University Bloomington) for sharing critical reagents, including the parental SINV infectious clone plasmids and E1/E2 polyclonal serum. We are grateful to Dr. Ram Savan (University of Washington) for sharing 5XISGF3-GLuc Huh7 reporter cells, and to Dr. Dan Stetson (University of Washington) for sharing *RIG-I KO, MDA5 KO,* and *MAVS* KO A549 cells. We thank the Hutch Shared Resources, especially the Genomics and Flow Cytometry Core Facilities, for technical support.

## Author contributions

Conceptualization: TB, TSF, EMA, HSM

Methodology: TB, TSF, EMA, FW, KM, LC, HSM

Investigation: TB, TSF, EMA, FW, LC

Formal Analysis: TB, TSF, EMA

Validation: TB, TSF, EMA

Visualization: TB, TSF, HSM

Resources: TB, TSF, KM, HSM

Data curation: TB, TSF, KM, HSM

Funding acquisition: TB, ME, KM, HSM

Project administration: TB, KM, HSM

Supervision: TB, ME, KM, HSM

Writing – original draft: TB, TSF, HSM

Writing – review & editing: TB, TSF, EMA, ME, KM, HSM

## Data and Materials Availability

All data needed to evaluate the conclusions in the paper are present in the paper and/or the Supplementary Materials.

## List of Supplementary Materials

Figures S1-S13, Data S1-S2

