## Supplemental Files for "A conserved in-frame stop codon acts as a multipotent defense mechanism in alphaviruses"

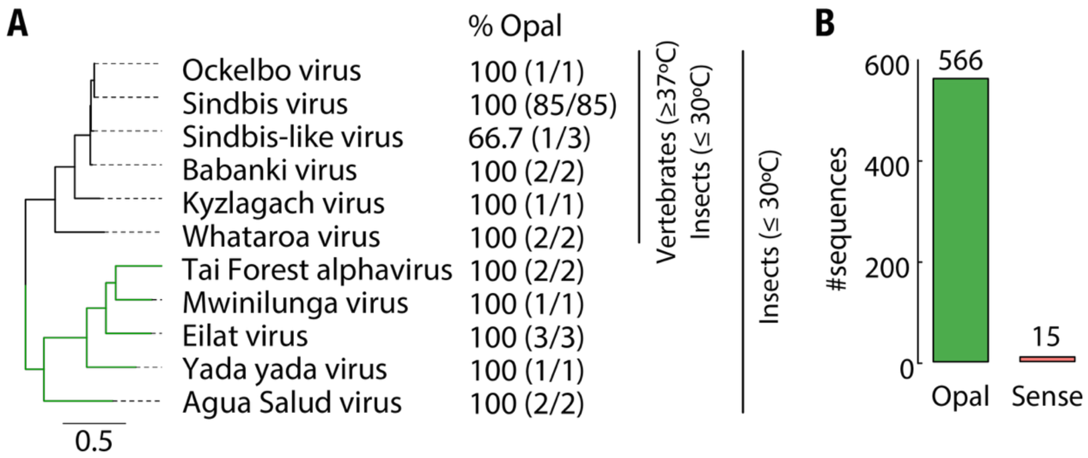


**Fig. S1. Conservation of the nsP3 opal stop codon among mosquito-isolated alphaviruses. (A)** A maximum-likelihood phylogenetic tree of two closely related dual-host and insect-specific alphavirus clades was built from whole-genome sequences. The percentage of occurrence of the nsP3 opal codon, shown next to each taxon, was calculated from publicly available sequences. However, except for Sindbis virus, which has 85 reported sequences, only a few sequences are available for each of the other viruses (shown in parentheses). **(B)** Percentage occurrence of the nsP3 opal codon and the extended PRT codon context UGAC among all available mosquito-isolated alphavirus sequences (n = 581). Numbers above each bar indicate the number of sequences analyzed.

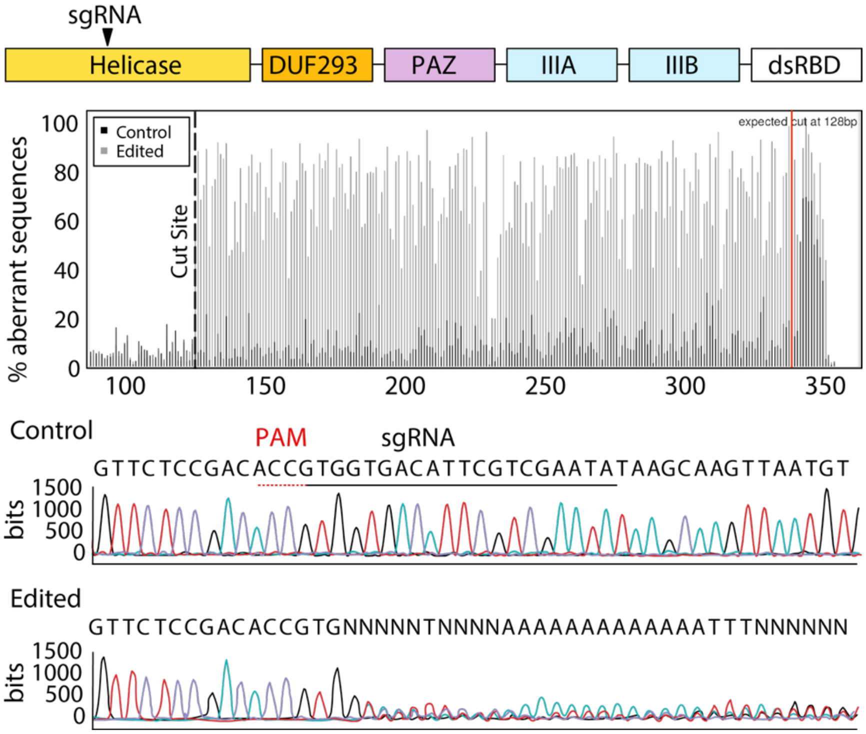


**Fig. S2. Editing efficiency in *Dcr2* KO U4.4 cells.** Domain structure of *the Aedes* Dicer 2 protein. The black arrow indicates the approximate location of the CRISPR sgRNA target. (Middle) TIDE analysis plot showing editing efficiency in U4.4 cells. The vertical dashed line marks the expected cut site. (Bottom) Chromatogram of the *Dcr-2* target region in control and CRISPR-edited cells. The region beyond the vertical red line shows a drop in sequence quality.

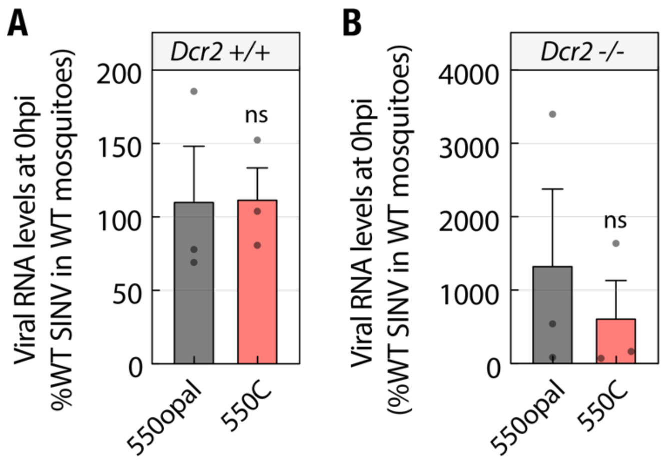


**Fig. S3. Initial viral inoculum levels in injected mosquitoes. (A)** Wild-type (*Dcr2 +/+*) and **(B)** *Dcr2* KO (*Dcr2 -/-*) *Aedes aegypti* mosquitoes were infected with SINV 550opal and 550C. Approximately 0-2 h after infection, injected mosquitoes were collected, and viral RNA was quantified by qRT-PCR. Data represent three independent biological replicates, each comprising five pooled mosquitoes. Error bars indicate the standard error of the mean (SEM). Unpaired t-test. ns = not significant.

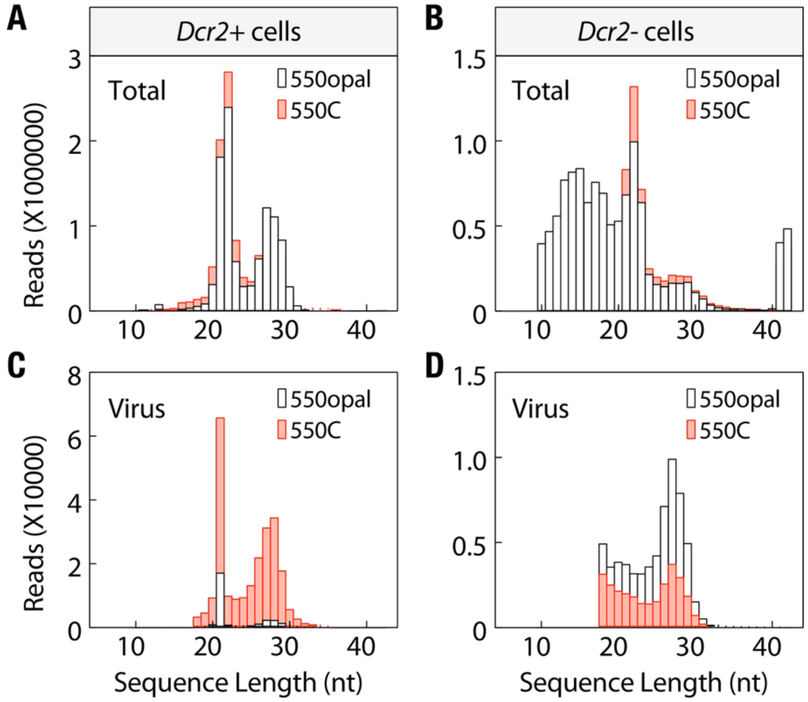


**Fig. S4. Small RNA read distribution in *Dcr2*+ U4.4 and *Dcr2*- C6/36 cells. (A-B)** Total small RNA read counts from *Dcr2*+ **(A)** and *Dcr2*- **(B)** cells infected with wild-type SINV 550opal and SINV 550C. **(C-D)** Virus-specific small RNA read counts from *Dcr2*+ **(C)** and *Dcr2*- **(D)** cells infected with wild-type SINV 550opal and SINV 550C. The data represent three independent biological replicates. Error bars indicate the standard error of the mean (SEM). Two-way ANOVA with Tukey's multiple-comparisons test. *** = $p < 0.001$; ns = not significant.

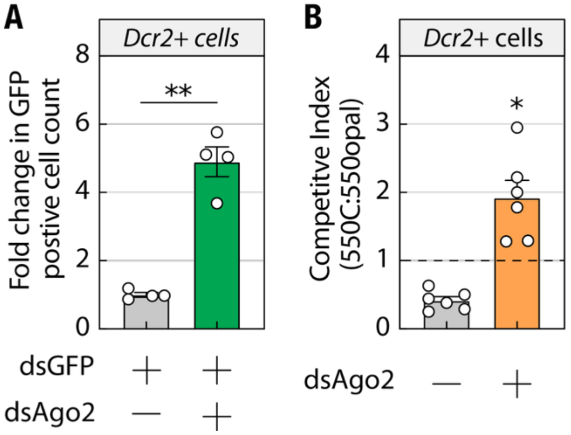


**Fig. S5. Effect of *Ago2* KD on siRNA activity and SINV 550C competitive fitness in *Dcr2*+ cells. (A)** Effect of dsRNA-mediated *Ago2* silencing on GFP reporter expression in *Dcr2*+ mosquito cells co-treated with *GFP* double-stranded RNA (dsGFP), as quantified by flow cytometry. Error bars represent the standard error of the mean (SEM). Student's t-test. ** = $p < 0.01$. Two-way ANOVA with Tukey's multiple comparisons test. **(B)** Competitive fitness of the SINV 550C variant against wild-type SINV 550opal in U4.4 cells treated with dsAgo2. The data represent five independent biological replicates. Error bars represent the standard error of the mean (SEM). One-sample t-test compared to a neutral competitive index (550C:550opal) of 1. **** = $p < 0.0001$, * = $p < 0.05$.

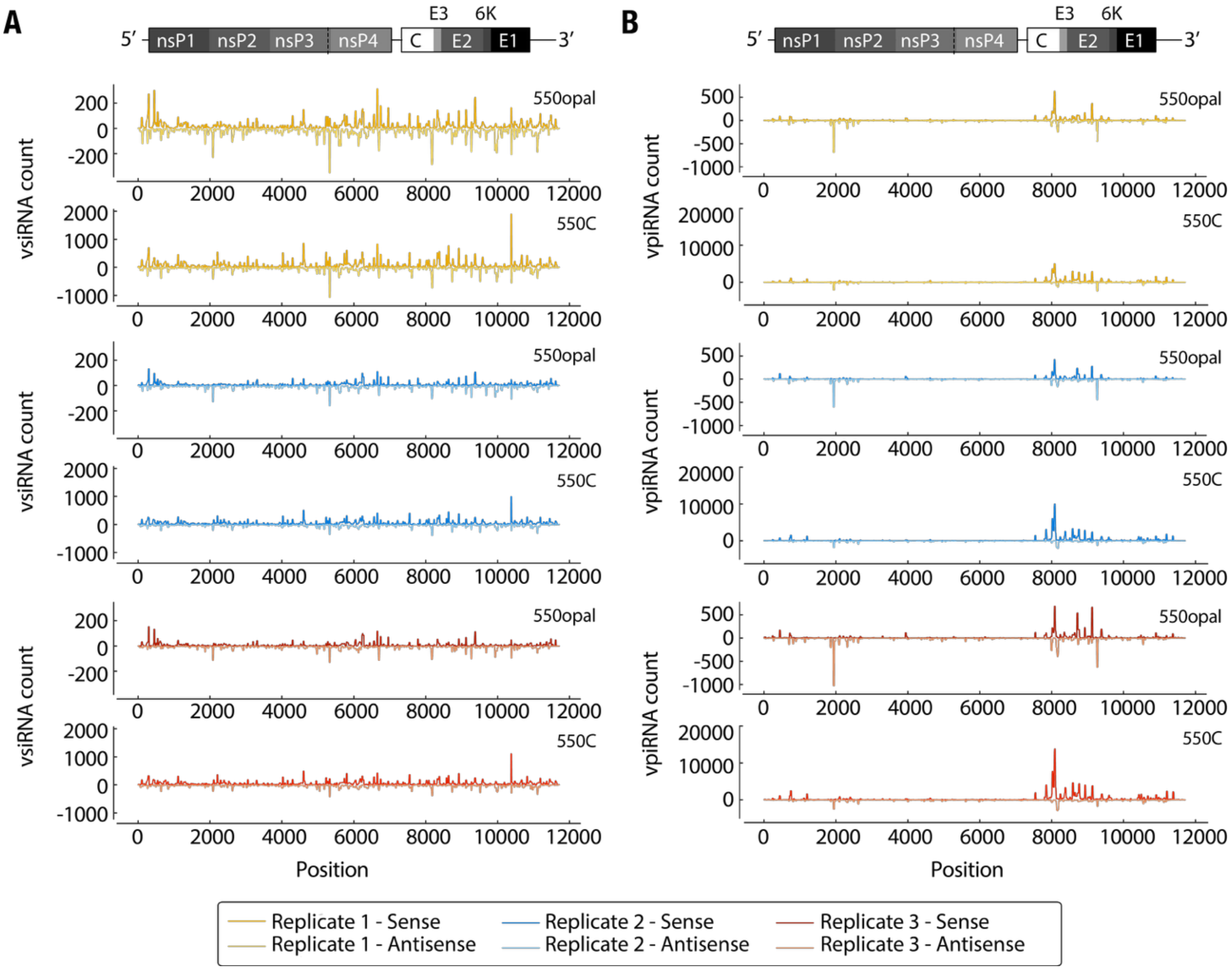


**Fig. S6. Replicate data for virus-mapped vsiRNA and vpiRNA in *Dcr2*+ mosquito cells. (A)** Normalized vsiRNA read distribution in *Dcr2*+ cells infected with wild-type SINV 550opal and SINV 550C. **(B)** Normalized vpiRNA read distribution in *Dcr2*+ cells infected with wild-type SINV 550opal and SINV 550C.

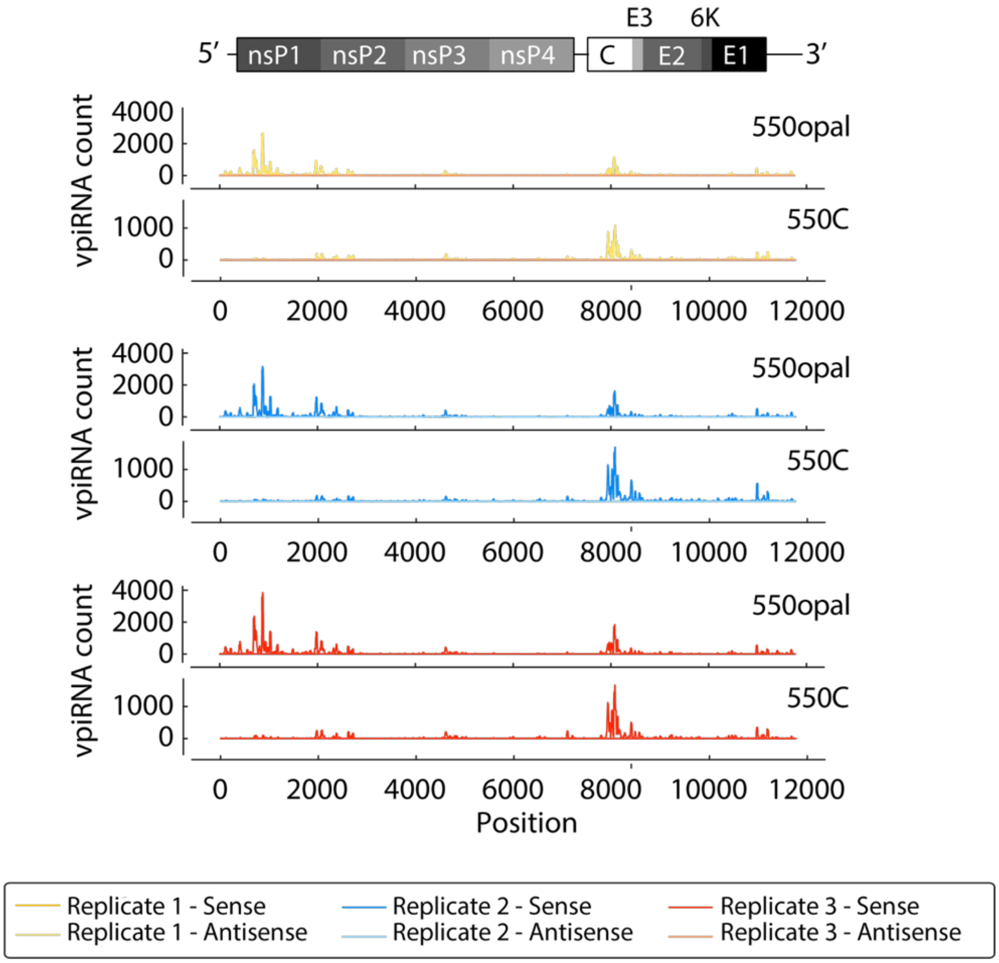


**Fig. S7. Replicate data for virus-mapped vpiRNAs in *Dcr2*– mosquito cells.** Normalized vpiRNA read distribution in *Dcr2*– cells infected with wild-type SINV 550opal and SINV 550C.

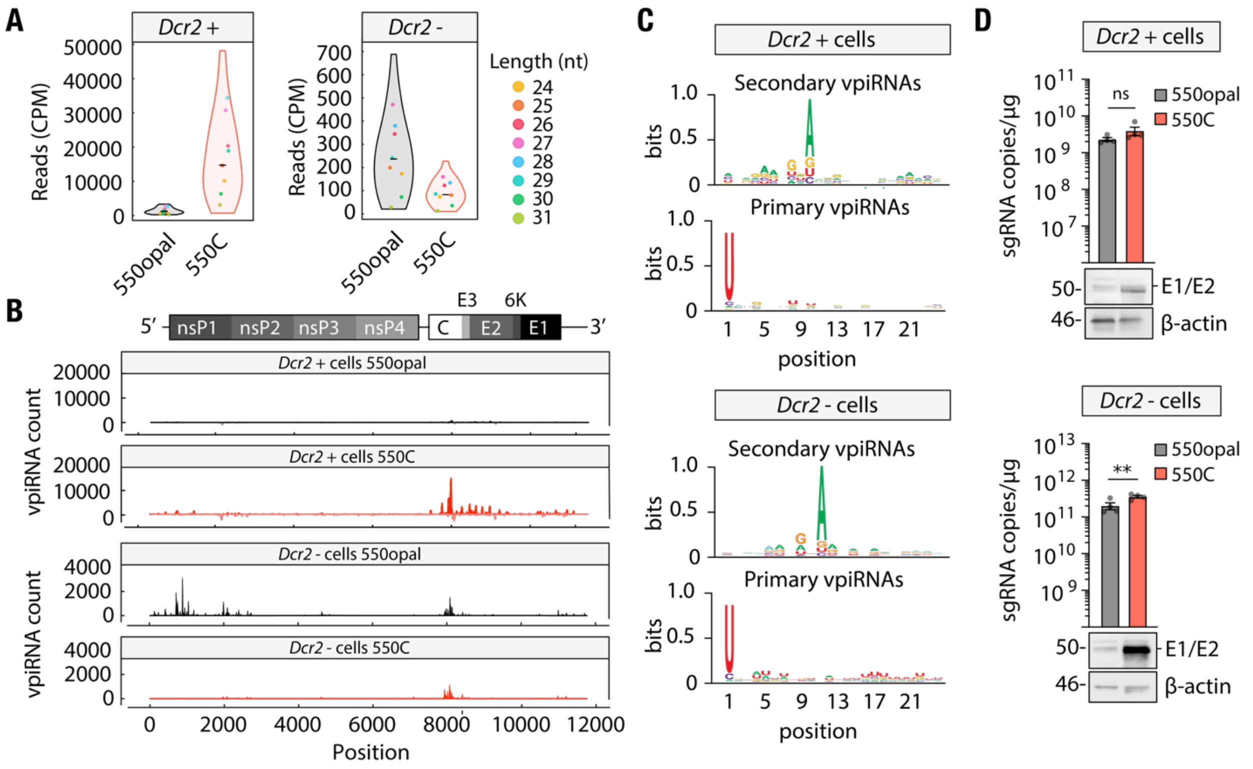


**Fig. S8. Virus-derived piRNAs are not antiviral in mosquito cells. (A)** Normalized median read counts of vpiRNAs across size classes in *Dcr2+* U4.4 cells infected with wild-type SINV 550opal or SINV 550C viruses. Student's t-test. **(B)** Distribution of vpiRNA reads from wild-type SINV 550opal (gray) or SINV 550C (red) viruses in *Dcr2+* U4.4 cells. Positive and negative Y-axis values indicate read counts mapped to the sense and antisense strands at each position along the SINV genome and subgenome (X-axis). **(C)** Cumulative per-position nucleotide frequency of the first 24 bases in sense (top) and antisense (bottom) vpiRNA reads from *Dcr2+* U4.4 cells infected with wild-type SINV 550opal or SINV 550C viruses. Logoplots were generated using WebLogo 3. **(D)** (top) Levels of intracellular viral subgenomic RNA in *Dcr2+* U4.4 cells infected with wild-type SINV 550opal or SINV 550C viruses, as quantified by qRT-PCR. (bottom) Western blot analysis of E1/E2 structural glycoprotein levels in *Dcr2+* U4.4 cells infected with wild-type SINV 550opal or SINV 550C viruses. The data represent four independent biological replicates. Error bars indicate the standard error of the mean (SEM). Student's t-test. **** = $p < 0.0001$, *** = $p < 0.001$, ** = $p < 0.01$, ns = not significant.

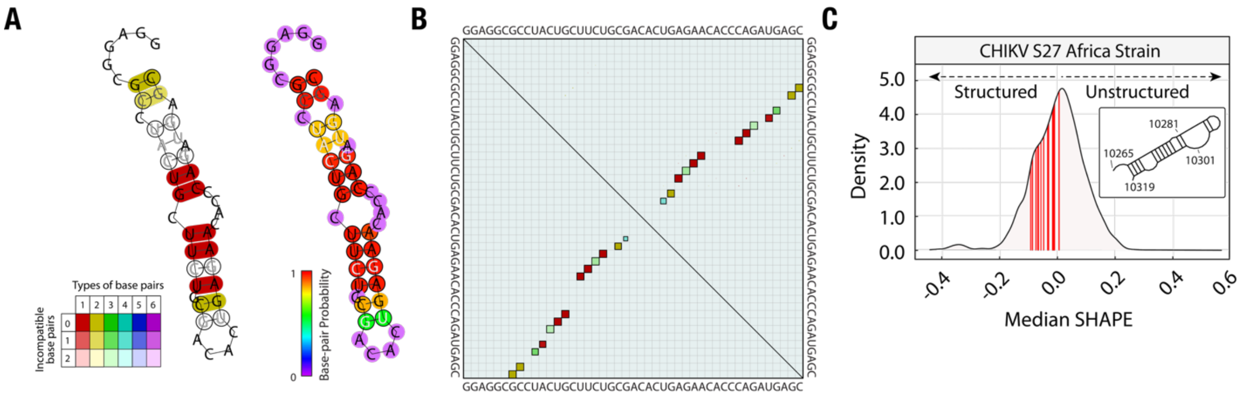


**Fig. S9. Sequence and structural conservation of E1-hs RNA across alphaviruses. (A)** (Left) RNAalifold-predicted structure of the E1-hs region spanning SINV nucleotides 10328-10372. Colors highlight the mutational pattern relative to the predicted RNA structure. The color key indicates whether a predicted base pair is formed by multiple nucleotide combinations or by consistent or compensatory mutations. Pale colors indicate the absence of base pairing in some sequences in the alignment. (Right) Base-pairing probability of the predicted RNA structure. **(B)** Base-pairing probability matrix of the E1-hs region. The upper triangle shows pairing probabilities, with box size proportional to the likelihood of a base pair (i, j). The lower triangle depicts the minimum free energy (MFE) structure using boxes of uniform size. In alignments, dot plots are color-coded to indicate sequence variation (same as panel A). Note: Some weakly conserved base pairs are not shown in panel A but are visible in panel B. **(C)** Density distribution profile of median SHAPE values of CHIKV RNA. Median SHAPE values for E1-hs residues are highlighted in red. The inset shows the SHAPE-constrained, RNAfold structure.

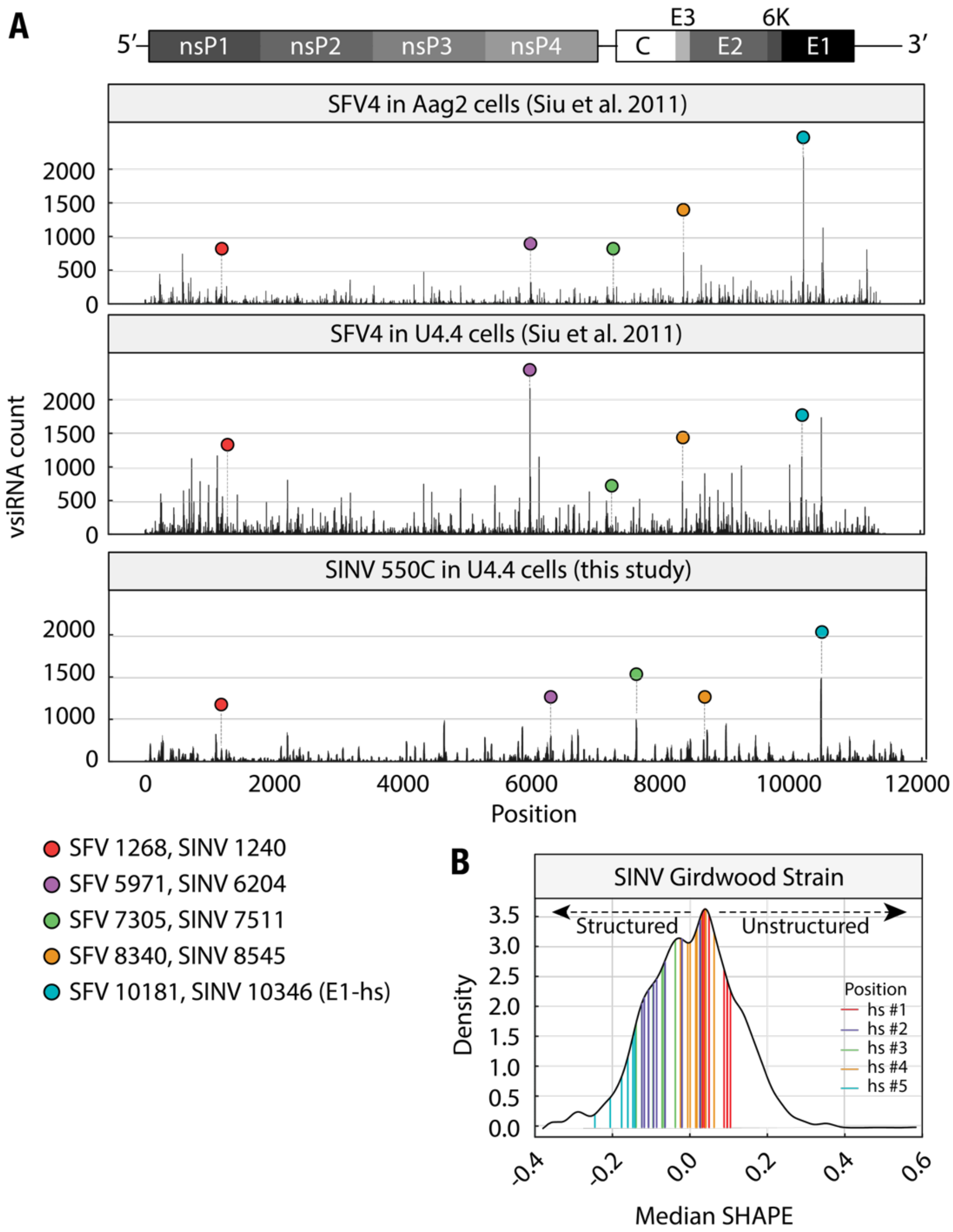


**Fig. S10. Shared vsiRNA hotspots between distantly related alphaviruses. (A)** vsiRNA hotspots identified in the Semliki Forest virus (SFV4) genome by *Siu et al.* in RNAi-competent *Aedes aegypti*-derived Aag2 (Top) and *Aedes albopictus*-derived U4.4 (Middle) cells. (Bottom) vsiRNA hotspots identified in SINV 550C in *Aedes albopictus*-derived U4.4 cells in this study. Four shared hotspots at SINV positions 1240, 7511, 8545, and 10346 are highlighted with colored circles (see key for details). **(B)** Density distribution profile of median SHAPE values for SINV RNA. Median SHAPE values for hotspot-region residues are highlighted in color.

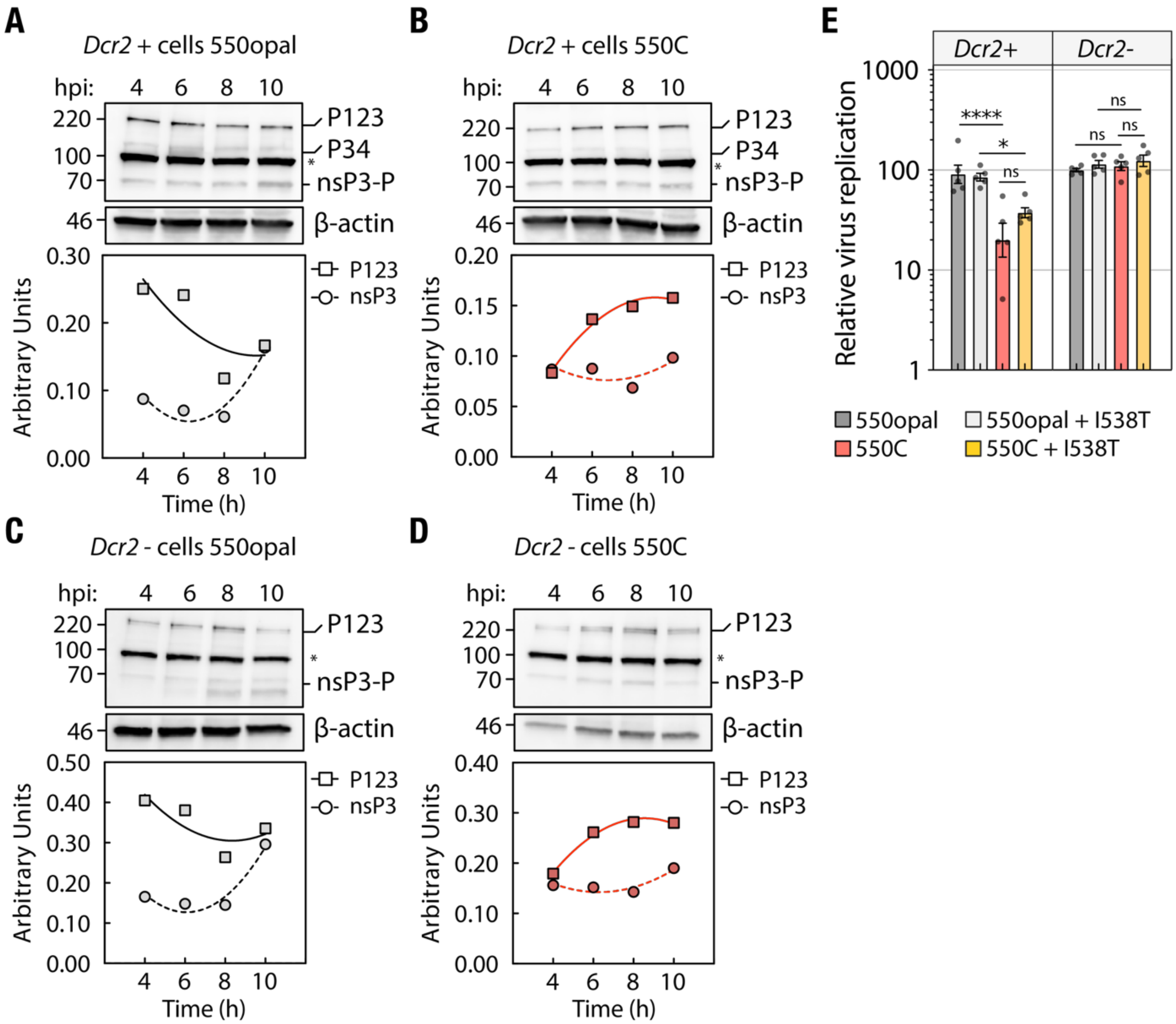


**Fig. S11. Non-structural polyprotein processing cadence in mosquito cells.** Temporal changes in the levels of unprocessed (P123) and processed (nsP3) nsP proteins, as determined by densitometric analysis of Western blot data from **(A-B)** *Dcr2+* (U4.4) and **(C-D)** *Dcr2–* (C6/36) cells infected with wild-type SINV 550opal **(A, C)** and SINV 550C **(B, D)**. Harvested protein lysates were probed with an anti-FLAG (nsP3) antibody. β-actin served as a loading control. A non-specific band cross-reactive with the anti-FLAG antibody is denoted with an asterisk (*). The data are representative of two independent experiments. **(E)** Replication of wildtype SINV 550opal or SINV 550C viruses with or without nsP1 I538T mutation in *Dcr2+* U4.4 cells (left) and *Dcr2–* C6/36 cells (right) as quantified via luciferase reporter assays at 48hpi. The data represent independent biological replicates (n=5). Error bars indicate the standard error of the mean (SEM).

Two-way ANOVA with Tukey's multiple comparisons test. **** = p < 0.0001, * = p < 0.05, ns = not significant.

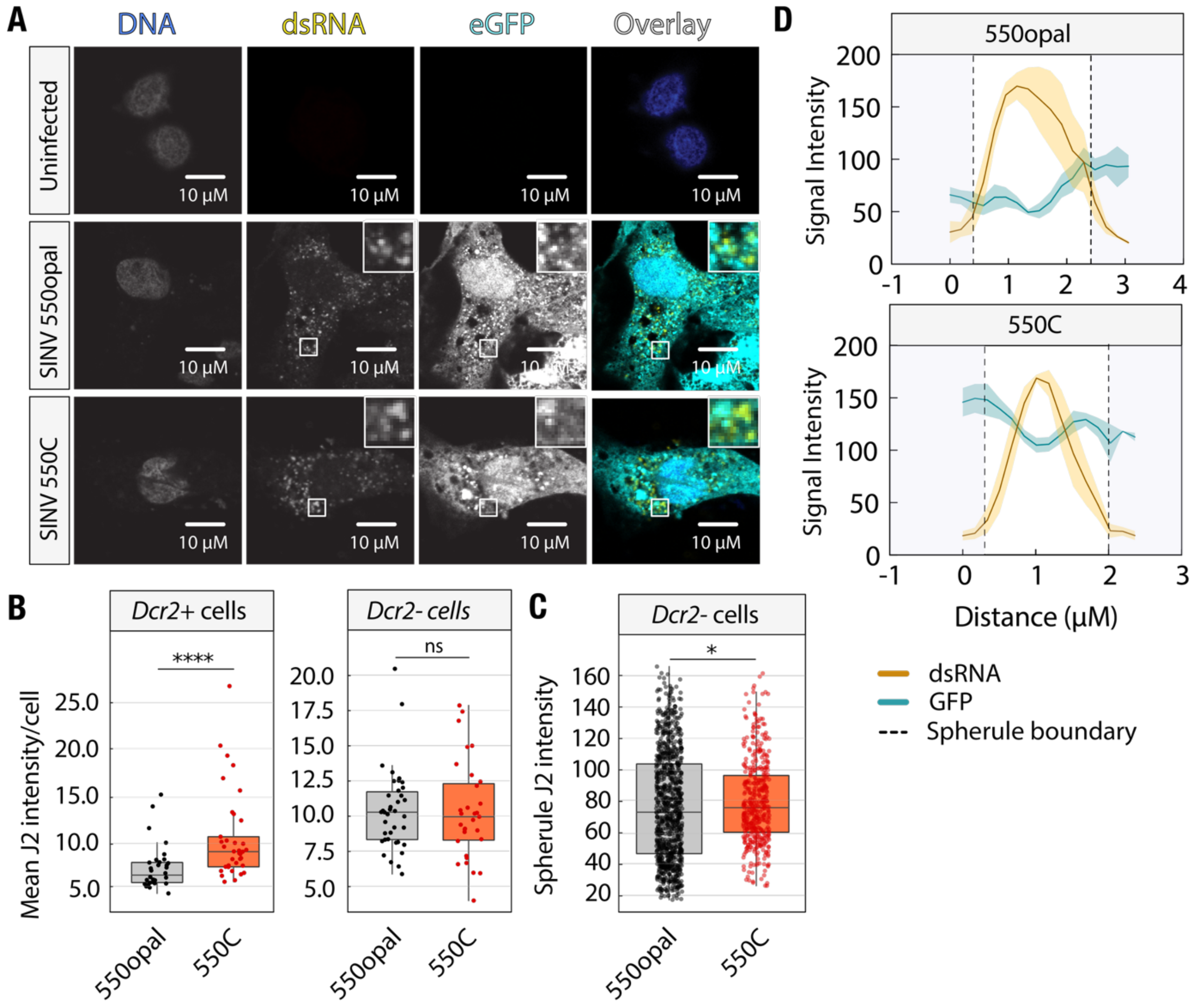


**Fig. S12. J2 signal quantification in SINV-infected *Dcr2-* cells. (A)** Localization of viral dsRNA (detected by the J2 antibody) and virally encoded GFP in *Dcr2-* C6/36 cells, either uninfected or infected with wild-type SINV 550opal (top) or SINV 550C (bottom), as determined by confocal microscopy. The inset shows GFP and dsRNA localization around viral replication spherules. **(B)** Mean J2 intensity per cell (n≥32) in *Dcr2+* U4.4 (left) and *Dcr2-* C6/36 (n≥31) (right) cells infected with wild-type SINV 550opal and SINV 550C. **(C)** Spherule J2 intensity in *Dcr2-* C6/36 cells (n≥435) infected with wild-type SINV 550opal and SINV 550C. Error bars indicate the standard error of the mean (SEM). Mann-Whitney U-test. **** = p < 0.0001, * = p < 0.05, ns = not significant. **(D)** Representative two-dimensional (2D) plot profile of spherules in *Dcr2+* cells (n=4) infected with wild-type and variant SINV. Dashed vertical lines indicate spherule boundaries.

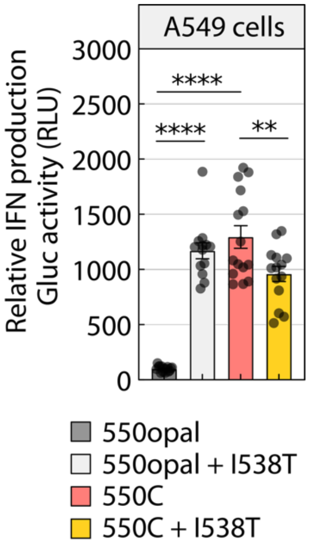


**Fig. S13. Effect of nsP1 mutation on SINV 550C-induced IFN response in human cells.** Reporter activity in 5XISGF3-GLuc Huh7 cells treated with supernatants collected from infected A549 cells 16 hours post-infection. The data represent the mean of n=14 biological replicates. Error bars indicate the standard error of the mean (SEM). One-way ANOVA with Tukey's multiple comparisons test. **** = $p < 0.0001$, ** = $p < 0.05$, ns = not significant.
